# What does the mean mean? A simple test for neuroscience

**DOI:** 10.1101/2021.11.28.469673

**Authors:** A. Tlaie, K. A. Shapcott, T. van der Plas, J. Rowland, R. Lees, J. Keeling, A. Packer, P. Tiesinga, M. L. Schölvinck, M. N. Havenith

## Abstract

Trial-averaged metrics, e.g. tuning curves or population response vectors, are a ubiquitous way of characterizing neuronal activity. But how relevant are such trial-averaged responses to neuronal computation itself? Here we present a simple test to estimate whether average responses reflect aspects of neuronal activity that contribute to neuronal processing. The test probes two assumptions implicitly made whenever average metrics are treated as meaningful representations of neuronal activity:

1. Reliability: Neuronal responses repeat consistently enough across trials that they convey a recognizable reflection of the average response to downstream regions.
2. Behavioural relevance: If a single-trial response is more similar to the average template, it is more likely to evoke correct behavioural responses.

We apply this test to two data sets: (1) Two-photon recordings in primary somatosensory cortices (S1 and S2) of mice trained to detect optogenetic stimulation in S1; and (2) Electrophysiological recordings from 71 brain areas in mice performing a contrast discrimination task. Under the highly controlled settings of data set 1, both assumptions were largely fulfilled. Moreover, better-matched single-trial responses predicted correct behaviour. In contrast, the less restrictive paradigm of data set 2 met neither assumption, with the match between single-trial and average responses being neither reliable nor predictive of behaviour. Simulations confirmed these results. We conclude that when behaviour is less tightly restricted, average responses do not seem particularly relevant to neuronal computation, potentially because information is encoded more dynamically. Most importantly, we encourage researchers to apply this simple test of computational relevance whenever using trial-averaged neuronal metrics, in order to gauge how representative cross-trial averages are in a given context.

## Introduction

Brain dynamics are commonly studied by recording neuronal activity over many stimulus repetitions (trials) and subsequently averaging them across time. Trial-averaging has been applied to single neurons, describing their average response preferences [1–6], and, more recently, to neural populations [7–9]. Implicit in the practice of trial averaging is the notion that deviations from the average response represent ‘noise’ of one form or another. The exact interpretation of such neuronal noise has been debated [10], ranging from truly random and meaningless activity [11–14], to neuronal processes that are meaningful but irrelevant for the neuronal computation at hand [15–17], to an intrinsic ingredient of efficient neuronal coding [18–20]. Nevertheless, in all of these cases a clear distinction is being made between neuronal activity that is directly related to the cognitive process under study (e.g. perceiving a specific stimulus) – which is approximated by a trial-averaged neuronal response – and ‘the rest’.

While this framework has undoubtedly been useful for characterizing the general response dynamics of neuronal networks, there is a sizable explanatory gap between the general neuronal response preferences reflected in trial-averaged metrics, and the way in which neurons transmit information moment-by-moment. As such, using trial-averaged data as a proxy to infer principles of one-shot, moment-by-moment neuronal processing is potentially problematic - an issue that has repeatedly been discussed in the field (see for instance [21–24]). However, neuroscience as a field has so far been reluctant to draw practical consequences. A vast majority of neuroscience studies present trial-averaged metrics like receptive fields, response preferences or peri-stimulus time histograms. These metrics rely on the implicit assumption that trial-averaged neuronal activity is fundamentally meaningful to our understanding of neuronal processing. For instance, upon finding that with repeated stimulus exposure, trial-averaged population responses become more sensitive to behaviourally relevant stimuli [3, 4], it is implicitly assumed that this average neuronal shift will improve an animal’s ability to perceive these stimuli correctly. In other words, neuroscience as a field seems to suffer from a disconnect between the limitations of cross-trial averaging that we acknowledge explicitly, and the implicit assumptions that we allow ourselves to make when we use cross-trial averages in our work.

One potential reason that this disconnect has not been tackled more actively is that the evidence regarding the functional relevance of trial-averaged responses is quite split. On the one hand, studies highlighting the large intertrial variability of neuronal responses [16, 17, 25–27] suggest that average responses fail to accurately capture ongoing neuronal dynamics. Then there is the simple fact that outside the lab, stimuli generally do not repeat, which renders pooled responses across stimulus repetitions a poor basis for neuronal coding. On the other hand, the fact that perceptual decisions can be altered by shifting neuronal activity away from the average response [28–31] indicates that at least in typical lab experiments [32], average population responses do matter [33]. Such widely diverging evidence suggests that cross-trial averages may be more relevant to neuronal computation in some contexts (and brain areas) than in others. This calls for a way to move the debate on their functional relevance beyond the realm of opinion and theory, and instead test this question concretely and practically across different experimental contexts.

In the present study, we provide a simple and widely applicable statistical test to explicitly determine whether cross-trial averages computed in a specific experiment are likely to be meaningful to neuronal information processing, or whether they are more likely to arise as an epiphenomenon with no clear computational function. To this end, our approach formalizes two implicit assumptions inherent in the computation of average neuronal responses, and tests directly whether they hold in a given experimental context (Fig. 1). Importantly, these two testable assumptions are not based on our own or other researchers’ views of how neuronal processing might actually work. Rather, they summarize how neuronal activity would need to behave if cross-trial averages reflect information that down-stream brain areas rely on to process information.

**FIG. 1.**
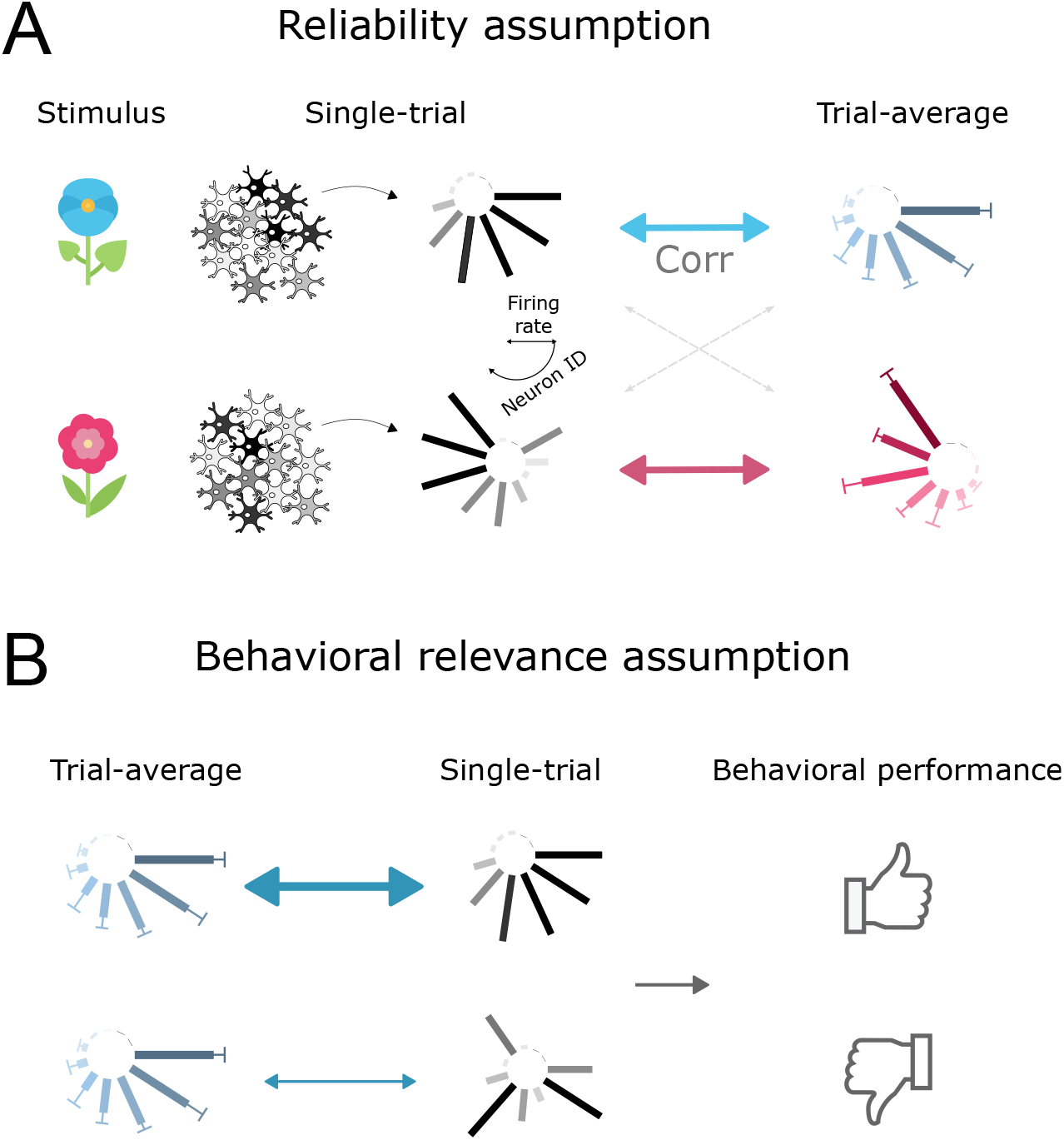
Two assumptions underlying the computation of average population responses. A) Reliability: single-trial responses correlate better with the trial-averaged response to the same stimulus, than with the trial-averaged response to a different stimulus. B) Behavioural relevance: better matched single-trial responses lead to more efficient behaviour.

1. Reliability: The responses of task-relevant neuronal populations repeat consistently enough to recognizably reflect the average response.The rationale for this assumption is that if neuronal responses varied so widely that the trial-averaged response was in no way recognizable from single-trial responses, then that would render the trial-averaged response uninformative to downstream neuronal processing.

2. Behavioural relevance: If a single-trial response better matches the average response, it is more successful in evoking correct behavioural responses. If cross-trial averages represent the ‘true signal’ of a neuronal population, which is obscured by single-trial noise, then the more similar a single-trial response is to the average, the higher its signal-to-noise ratio; therefore, the easier the information readout for downstream areas; therefore, the better the chances of a successful behavioural response.

To quantify to what extent a given data set adheres to each of these assumptions, we developed two simple statistical indices, and benchmarked them on two complementary data sets featuring neuronal recordings in behaving mice.

## Results

We started by examining our two assumptions in a data set that was acquired under extremely tightly controlled experimental settings. Data set 1 consists of two-photon calcium imaging recordings in primary and secondary somatosensory cortex (S1 and S2) as mice detected a low intensity optogenetic stimulus in S1 [34] (Fig 2A). Mice were trained to lick for reward in response to the optogenetic activation of 5 to 150 randomly selected S1 neurons (‘stimulus present’ condition). On 33% of trials, there was a sham stimulus during which no optogenetic stimulation was given (‘stimulus absent’ condition). Simultaneously, using gCAMP6s, 250 − 631 neurons were imaged in S1 and 45 − 288 in S2. Notably, in S1, the stimulus directly drives the neuronal response, skipping upstream neuronal relays.

**FIG. 2.**
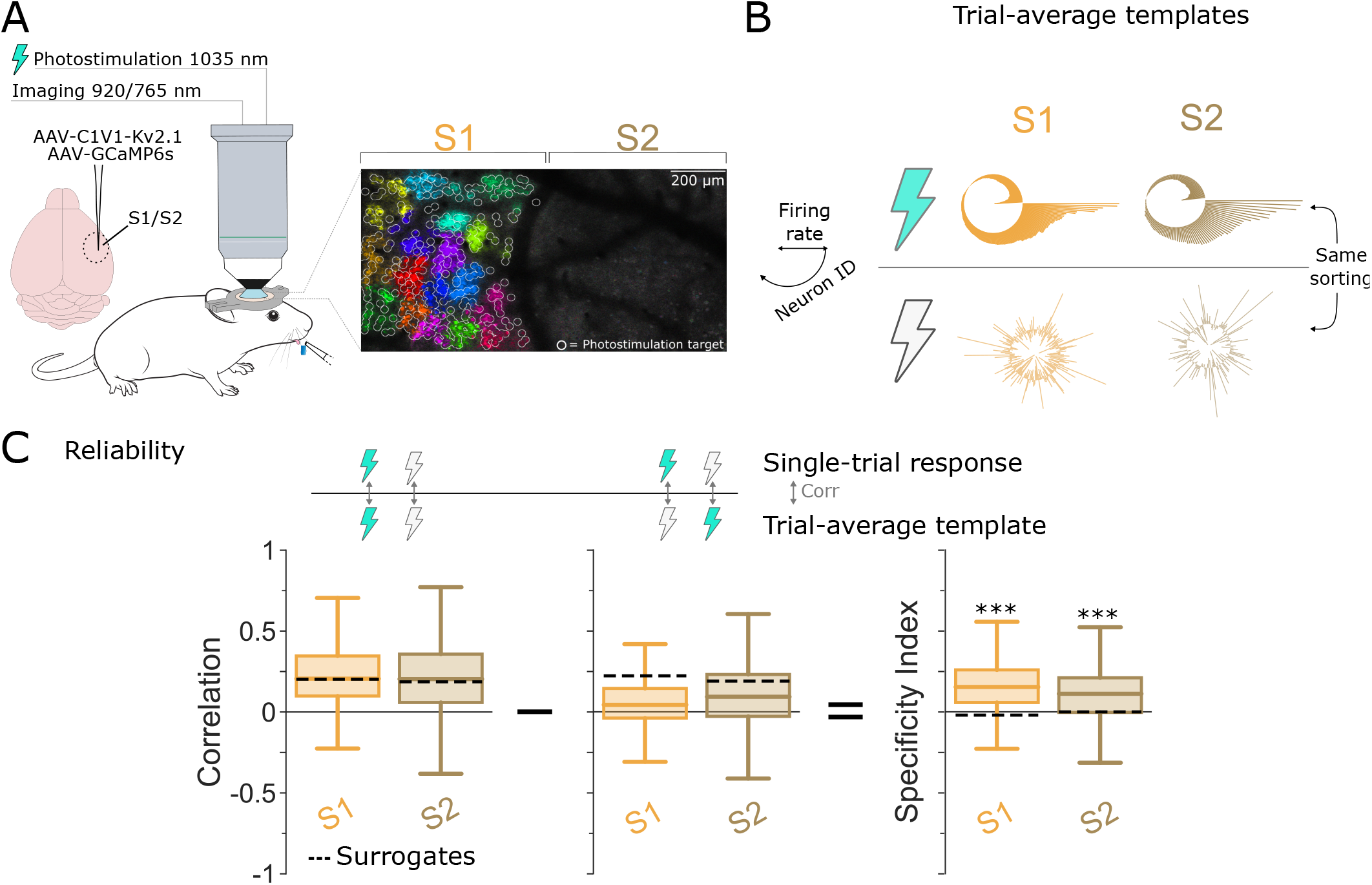
Single-trial responses are stimulus-specific for Data set 1. A) Animals report whether they perceived the optogenetic stimulation of somatosensory neurons (S1) through licking to receive reward. This panel was adapted from [34] with permission from the authors. B) Trial-average population responses (‘templates’) for S1 (orange) and S2 (brown), under optogenetic stimulation (top) or no stimulation (bottom) conditions. Neurons are sorted the same under both conditions. C) Distribution of the correlations between single-trial responses and the matching (left) and non-matching (middle) trial-averaged response templates. Box: 25^*th*^ and 75^*th*^ percentile. Center line: median. Whiskers: 10^*th*^ and 90^*th*^ percentile. Dotted lines: median of bootstrapped data. The difference between the correlations to the matching and non-matching templates gives the Specificity Index (right).

To probe the computational role of averages within this tightly controlled setting, we first computed average population responses for the two experimental conditions. Since individual stimulation intensities were often only presented in a small number of trials, we pooled all stimulation intensities (see Methods) into the ‘stimulus present’ condition (the high correlations between the average responses to different stimulation intensities are shown in Fig. S2A).

Average responses *templates* were computed as the mean fluorescence (Δ*F/F*) of each neuron in a time window of 0.5 s following the stimulation offset (Fig. 2B). We next quantified how well single-trial responses matched the corresponding average template (stimulation present or absent) (Fig. 2C, left; see also [35]). To this end, we computed linear correlations between the single-trial and trial-averaged population responses. While in principle, single-trial responses could reflect the corresponding average template in a multitude of ways, including multi-dimensional and/or non-linear relations, linear correlations are the correct way to capture their match if one accepts the assumptions underlying cross-trial averaging. Averages are based on linear computations (sums and rescaling), which implicitly assumes that the single-trial responses subsumed in the average differ from each other linearly and along one dimension - otherwise, pooling them in a linear average would not be a suitable approach.

Single-trial correlations were mostly positive in both S1 and S2 (Fig. S2C) (*n* = 1795 trials; *p <* 0.001), suggesting that single-trial responses represented the template quite faithfully. To assess if single-trial responses can be regarded as down-sampled or ‘noisy’ versions of the average template, we repeatedly computed a bootstrapped response vector for each trial based on the neurons’ average response preferences (Fig. S4, Methods). The surrogate data correlated only slightly better to the template than the original data in both S1 and S2 (Fig. 2C), suggesting that in Data set 1, single-trial responses can be interpreted as mostly faithful samples of the respective average template.

Correlations between single-trial responses and the population template may partially stem from neurons’ basic firing properties, which would not be task-related. To estimate the stimulus specificity of the correlations we observed, we also computed single-trial correlations to the incorrect template (e.g. ‘stimulus absent’ for a trial featuring optogenetic stimulation). Correlations to the incorrect template were significantly lower than to the correct one (Fig. 2C, middle, Mann-Whitney U-test, *p* = 5.98 * 10^−169^, *p* = 4.86 * 10^−51^ for S1 and S2, respectively). To quantify this difference directly, we defined the *Specificity Index*, which measures, on a single-trial basis, the excess correlation to the correct template compared to the incorrect template. Thus, the Specificity Index quantifies to what extent neuronal activity in an individual trial relates to the average response of the relevant experimental condition, compared to the average responses for other experimental conditions. Since the Specificity Index subtracts two correlation coefficients from each other, it is bounded between -2 and 2. For this data set, the Specificity Indices of single-trial responses indicate clear stimulus-specificity (Fig 2C, right). Note that while single-trial correlations to the average template scale directly with the amount of inter-trial variability in a data set, the Specificity Index is largely independent of inter-trial variability, because it reflects the differential match of single-trial responses to the correct versus incorrect template (see Fig. S6). Together, these results indicate that single-trial responses in Data set 1 were strongly and selectively correlated to the corresponding average template, largely fulfilling Assumption 1.

Next, we set out to test if the correlation between single-trial responses and average templates predicted the animal’s licking behaviour (see Fig 3A, left, for an example session). To this end, we separately examined the single-trial correlations in trials that resulted in hits, misses, correct rejections (CRs) and false positives (FPs) (Fig 3A, right). For the trials where optogenetic stimulation was present, single-trial correlations in S1 were significantly higher in hit trials than in miss trials, suggesting that a better match to the average template did indeed produce hit trials more often (Fig. 3B, Mann-Whitney U-test, *p* = 4.96*e* − 68). Similarly, while single-trial correlations were overall lower in the absence of optogenetic stimulation, correct rejections nevertheless featured significantly higher correlations than false positives (Fig. 3B, Mann-Whitney U-test, *p* = 2.82*e* − 17 for the hit/miss comparison, for each area). The same pattern held true for S2, though overall correlations were marginally smaller and the difference between correct and incorrect trials was somewhat less pronounced (Fig 3B; *p* = [2.02*e* − 50, 1.37*e* − 11] for hit/miss and CR/FP comparisons, respectively). To quantify directly to what extent single-trial correlations predicted behaviour, we computed the *Behavioural Relevance Index* (Ω) as Ω = *max*(*A*, 1 − *A*), where *A* is the Vargha-Delaney’s effect size [36] (Methods). The Behavioural Relevance Index quantifies whether successful behavioural responses occur preferentially in trials with a higher Specificity Index. The Behavioural Relevance Index is bounded between 0.5 and 1, with 0.5 indicating complete overlap between the distributions of Specificity Indices for correct and incorrect trials, and 1 meaning no overlap at all. For both the trials with stimulation (hits and misses) and without stimulation (CRs and FPs) Ω exceeded 0.5 in S1 and S2 (Fig. 3C). This suggests that in both areas, single-trial responses that were better matched to the cross-trial average resulted in more successful behaviour, fulfilling Assumption 2. Together, these results indicate that in Data set 1, cross-trial averages are both reliable and behaviourally relevant enough to serve a computationally meaningful function.

**FIG. 3.**
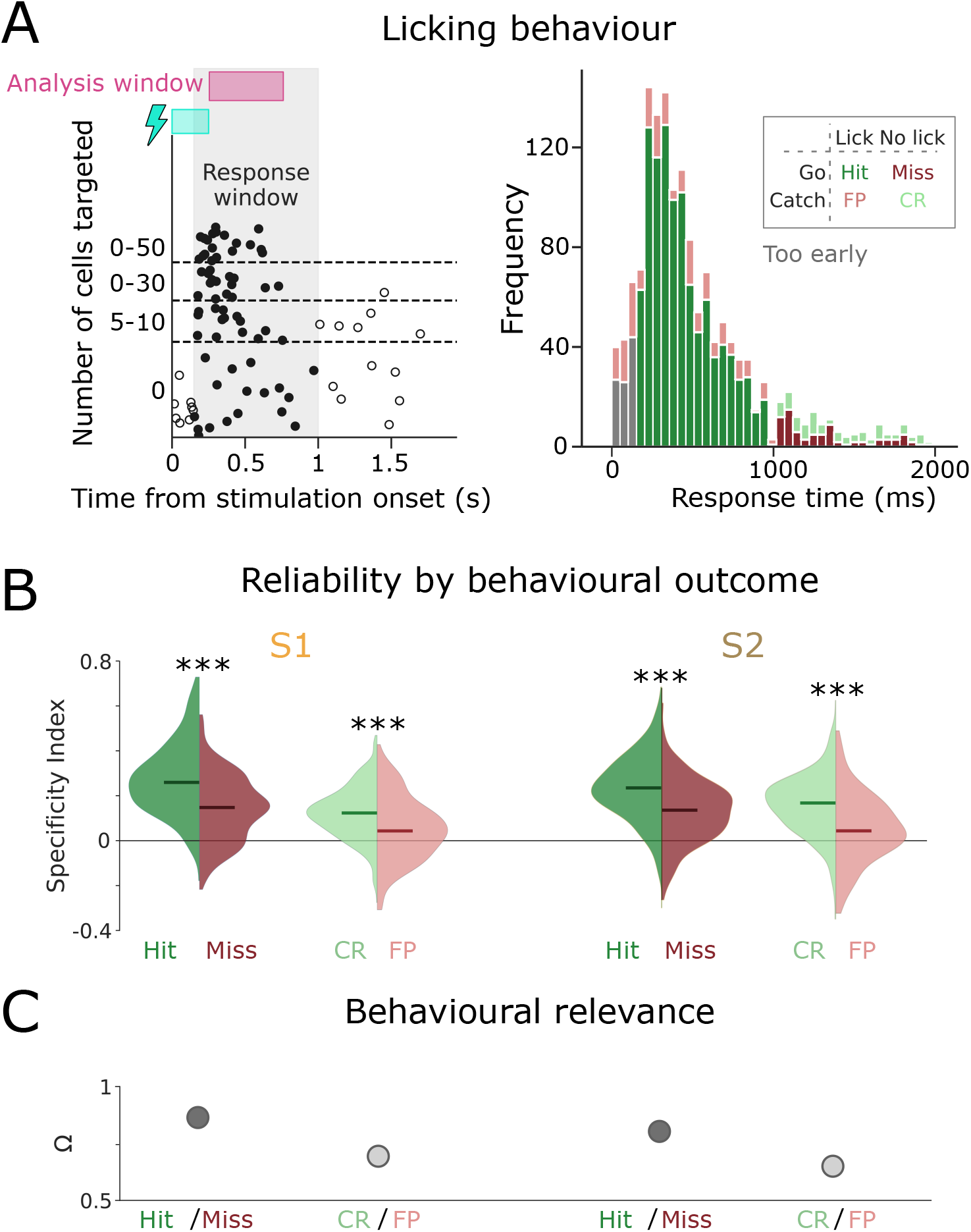
Better template-matching predicts better behaviour. A) Licking times for an example session (left) and for all sessions (right). The stimulation window is shown in blue, the analysis window in pink. B) Reliability of single-trial responses, as quantified by the Specificity Index, split out by hits, misses, correct rejections and false positives. C) Behavioural Relevance indices for these categories.

Building on these results, we set out to determine how computationally meaningful cross-trial averages might be within the less restrictive experimental paradigm of Data set 2. Data set 2 contains high-density electrophysiological (Neuropixel) recordings across 71 brain regions (Fig. 4A, right) in mice performing a two-choice contrast discrimination task [31]. Mice were presented with two gratings of varying contrast (0, 25, 50 or 100%) appearing in their left and right hemifield. To receive reward, animals turned a small steering wheel to bring the higher-contrast grating into the center, or refrained from moving the wheel if no grating appeared on either side (Fig. 4A, left). When both stimulus contrasts were equal, animals were randomly rewarded for turning right or left. Those trials were discarded in our analysis since there is no ‘correct’ behavioural response.

**FIG. 4.**
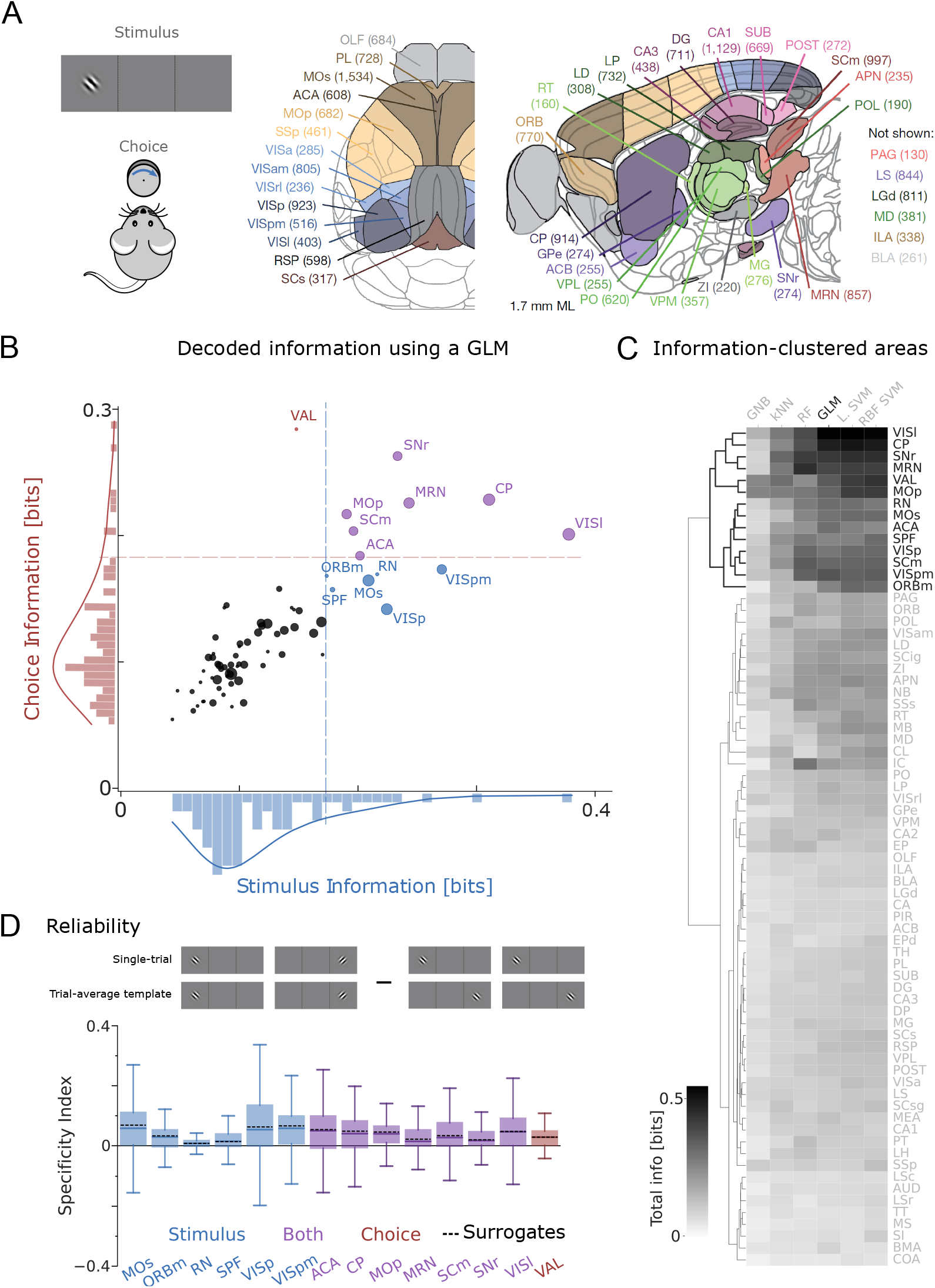
Single-trial responses are hardly stimulus-specific for Data set 2. A) Animals move a steering wheel to bring the higher-contrast grating stimulus into the centre (left), while being recorded from 71 brain areas (right). B) Stimulus and target choice information decoded by a multinomial GLM decoder (Methods) from the neuronal activity in all recorded brain areas. Each point represents the median (dot location) and standard deviation across sessions (dot size) of one brain area (see in-figure labels). Colours (blue, red, purple) represent those areas where (stimulus, choice, both) information was above an elbow criterion. C) We repeated the decoding with other models (see labels) and then performed a hierarchical clustering of the total mutual information of the ranked brain areas (rows). The 14 areas we found with the GLM (see B) are consistently found with other decoders. D) Specificity Index of the selected areas, defined as the difference in the correlations between single-trial responses and the matching (cartoon, left) and non-matching (cartoon, right) trial-averaged response templates. Box: 25^*th*^ and 75^*th*^ percentile. Center line: median. Whiskers: 10^*th*^ and 90^*th*^ percentile. Shaded areas: 5^*th*^ and 95^*th*^ percentiles of bootstrapped data. Dotted lines (surrogates): medians of the bootstrapped values for each recorded area.

Since this data set contains neuronal recordings from 71 brain areas, not all of which may be directly involved in the perceptual task at hand, we used a data-driven approach to identify to what extent neuronal population activity predicted the presented stimulus and/or the animal’s target choice. We trained a decoder (Multinomial GLM, Methods) based on single-trial population vectors, to identify either target choice (left/right/no turn) or stimulus condition (higher contrast on left/right, zero contrast on both). For the neuronal response vectors, we considered neuronal activity 0 − 200*ms* post-stimulus onset (Fig. S5). We then computed the mutual information between the decoder predictions and the real outcomes (Fig. 4B; Methods).

Many brain areas appeared to contain little task-relevant information (shown in black in Fig. 4B). We therefore used an elbow criterion (Methods) to determine a threshold for selecting brain areas that provided the highest information on either stimulus 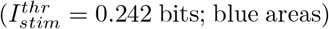, choice 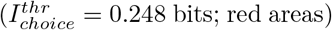, or both (i.e. both thresholds exceeded; purple areas). These areas seem largely congruent with the literature. For instance, primary visual cortex (VISp) is expected to reflect the visual stimulus, while choice information is conveyed e.g. by the ventral anteriorlateral complex of the thalamus (VAL) – known to be a central integrative center for motor control [37]. As an example of a both choice- and stimulus-informative area, we see caudoputamen (CP) - involved in goal-directed behaviours [38], spatial learning [39], and orienting saccadic eye movements based on reward expectancy [40]. To further validate our selection of relevant brain areas, we repeated the analysis with five other decoders (Fig. 4C). We ranked the total amount of Mutual Information per area (stimulus + choice information), for each of these models. Finally, we performed a hierarchical clustering and show that the areas we identified in Fig. 4B consistently appeared in the top performing cluster (Fig. 4C, highlighted in black). Together, these results converged on a group of 14 brain areas that conveyed significant task information regardless of the decoder approach.

Having identified task-relevant brain areas, we used neuronal recordings from those informative areas to test the two assumptions set out in the Introduction. First, we computed the average population response templates for different experimental conditions. To avoid working with trial numbers as low as *n* = 2 for specific contrast combinations, we pooled several contrast levels requiring the same behavioural response (e.g. 50% right – 0% left and 100% right – 50% left) into two conditions: Target stimulus on the left or on the right. Average responses to the individual contrast levels were very comparable (Fig. S2).

To test the first assumption, as we did for Data set 1, we quantified how well single-trial responses correlated with the template for a given stimulus (Fig. S3 A; see also [35]). Median correlations ranged from *r* = 0.56 to 0.89 across all brain areas (*n* = 89 to 3560 trials per brain area; all *p <* 0.001), suggesting that single-trial responses clearly resembled the average template. As a control, we computed 100 bootstrapped response vectors for each trial by drawing the same number of spikes as the original data, but based on each neuron’s average spiking probability (Fig. S3 B; Methods). These surrogate data uniformly correlated better to the template than the original data (Fig. S3 B), indicating that in Data set 2, single-trial responses exhibited more variation than explained by (Poissonian) down-sampling. As in Data set 1, single-trial correlations scaled with the amount of inter-trial variability, reflected for instance by the Fano Factor (S5). Note that the Fano Factors for all analysed brain areas in Data Set 2 fell within the range of previously reported results [41, 42] (Fig. S5 B), suggesting that Data set 2 provides a representative benchmark for neuronal activity in these brain areas.

Next, we estimated the stimulus specificity of the observed correlations by computing single-trial correlations to the incorrect template (e.g. ‘target right’ for the left target; see Fig 4 D, top). These were widely distributed, but on average marginally lower than the single-trial correlations to the correct template (Fig. S3B). Consequently, the Specificity Indices were mostly positive but rarely exceeded 0.1 (Fig. 4D, bottom). In other words, correlations between single-trial responses and template were largely stimulus-independent. This lack of response specificity was not directly predicted by the amount of inter-trial variability, because the Specificity Index reflects the differential link to correct versus incorrect template rather than the overall reproducibility of neuronal responses (see Fig. S5 B).

These results tally with recent work demonstrating how strongly non-task-related factors drive neuronal responses even in primary sensory areas like visual cortex [16, 17, 43–47]. However, the animal still needs to arrive at a coherent perceptual choice (e.g. steering right or left, see Fig. 5A). To test if trial-averaged templates are relevant to this perceptual decision, we compared single-trial correlations for hit trials (correct target choice) and miss trials (incorrect target or no response). Single-trial correlations were marginally lower in miss trials than in hit trials across most brain areas (Fig. 5B; Supp. Results). However, their difference was small, leading to Behavioural Relevance indices between 0.51 and 0.66. According to the Vargha & Delaney’s effect size, such values would be considered largely negligible, indicating that single-trial correlations are not a reliable way to predict subsequent behaviour in Data set 2 (Ω; Fig. 5 C; see also [36]).

**FIG. 5.**
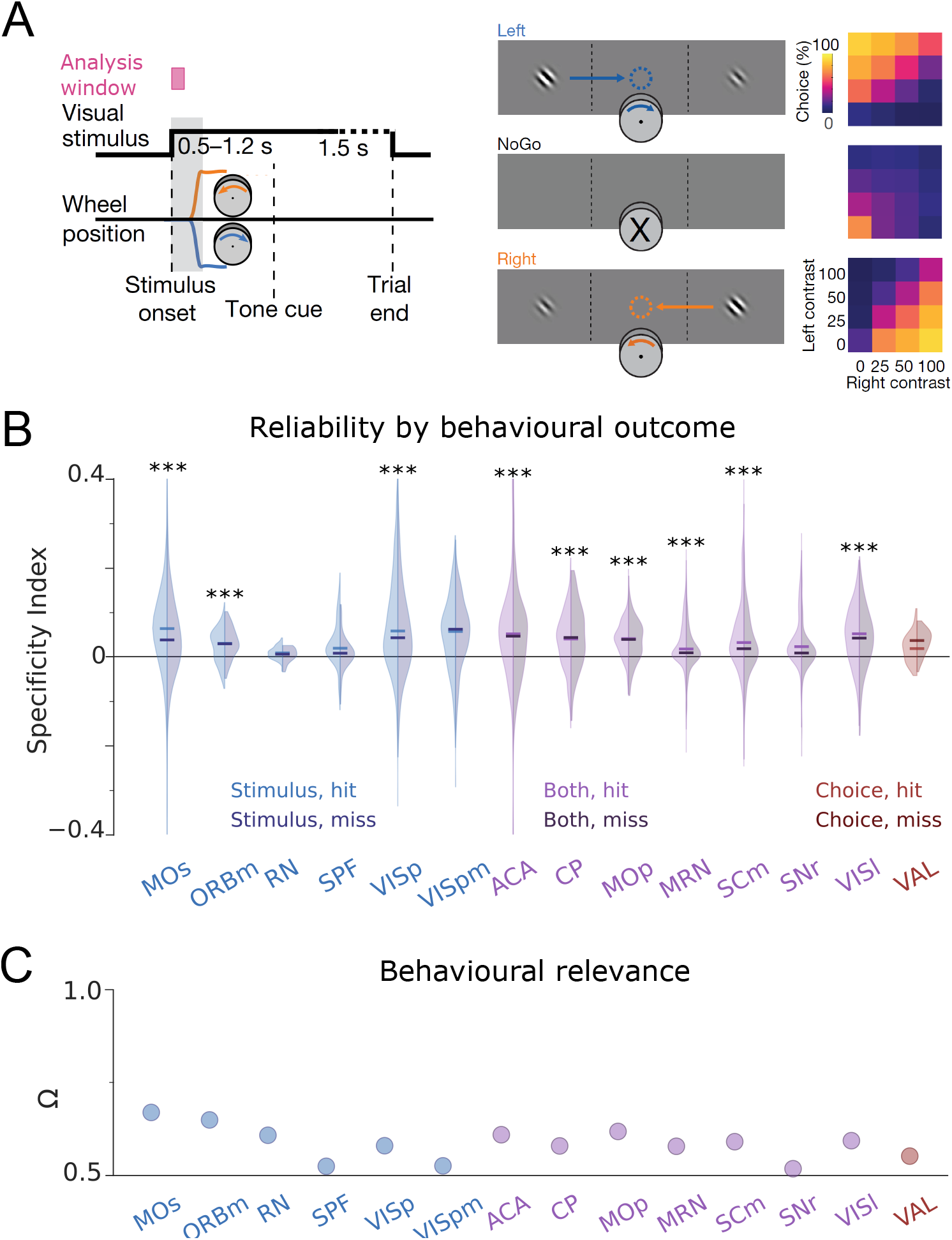
Single-trial responses are barely behaviourally relevant for Data set 2. A) Task. Mice bring the higher contrast stimulus to the centre by steering a wheel, or refrain from moving the wheel when no stimulus is present (left, middle) with high proficiency (right). B) Specificity index for the selected areas, split by hits and misses. C) Behavioural relevance for selected brain areas.

Together, these results suggest that in Data set 2, the relation between single-trial responses and trial-averaged templates is neither reliable nor specific, and does not appear to substantially inform subsequent behavioural choices. However, this estimate may present a lower bound for several reasons. First, while the task information conveyed by cross-trial averages seemed to be limited in the recorded population of neurons, it might be sufficient to generate accurate behaviour when scaled up to a larger population. To explore this possibility, we sub-sampled the population of recorded neurons in each brain area from *N/*10 to *N* . We then extrapolated how the Specificity and Behavioural Relevance Indices would evolve with a growing number of neurons. These extrapolations indicated that average template matching is unlikely to become more consistent, specific or behaviourally relevant with more neurons (Fig. 6 A, right). This did not seem to be a general feature of our extrapolation approach: for Data set 1, the Specificity Index appeared to remain largely stable with growing *n*, but Ω rose steeply, indicating that with a larger number of neurons, the match of individual trials to the average template would more strongly predict behaviour (Fig. 6 A, left).

**FIG. 6.**
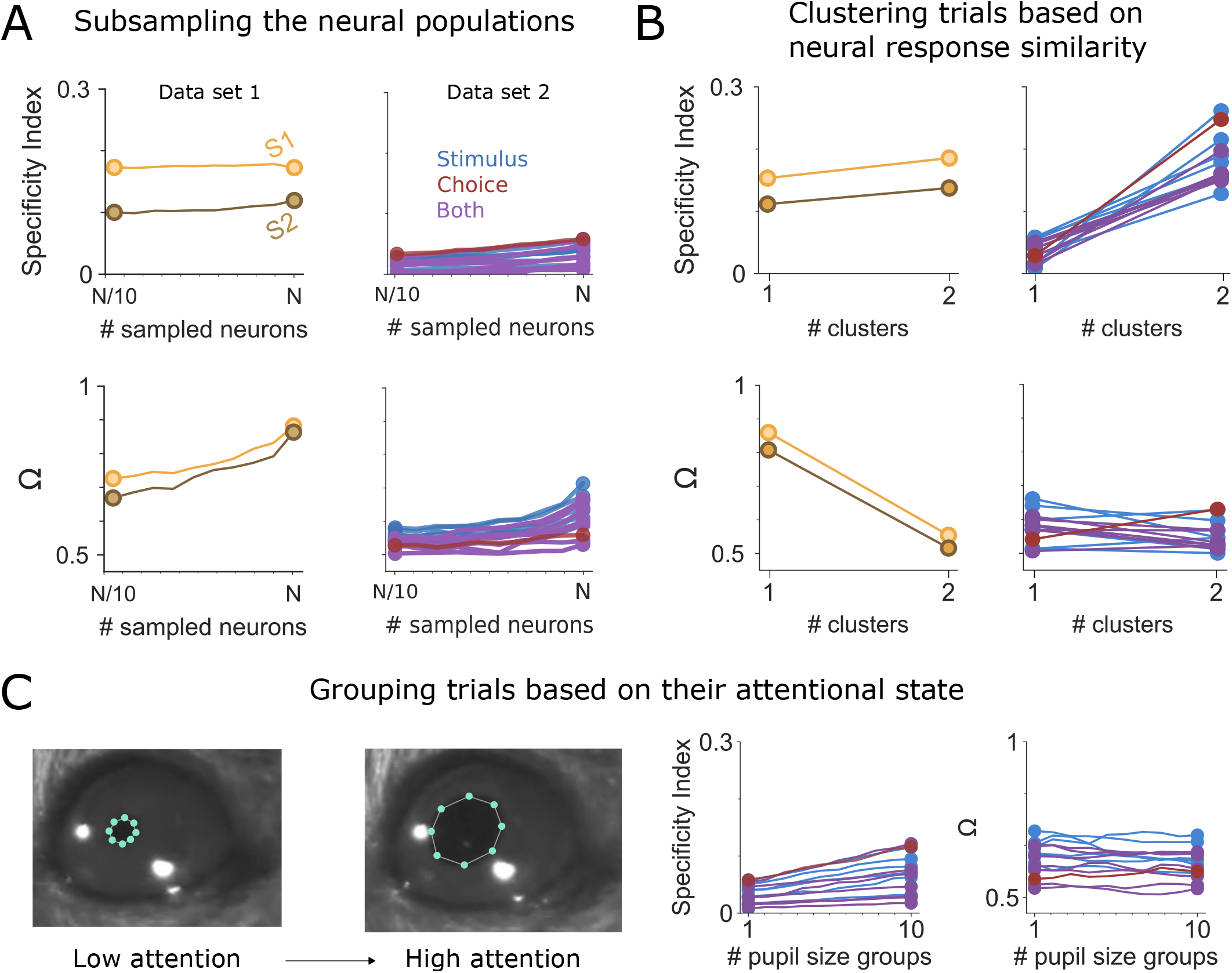
Control analyses for both data sets. A) We subsampled the neuronal populations to check whether we could extrapolate a marked benefit from adding neurons when performing template-matching. Increasing the number of sampled neurons left the Specificity Indices for both Data sets largely unchanged (top), and yielded slight increases in Ω (bottom). We then clustered trials based on the similarity in their neural response (B) and pupil size (C). Neither of these clusterings made template-matching more stimulus-specific or behaviorally relevant.

Alternatively, there may be a ‘supergroup’ of highly reliable neurons, which might predominantly drive downstream processing [48, 49]. However, a jackknife procedure (i.e. removing one neuron at a time from the data; see Methods) revealed no neurons that particularly boosted single-trial correlations, the Specificity Index, or the Behavioural Relevance Index (Fig. S9). In some areas (e.g. RN or SNr), there seemed to be at least some neurons that contributed substantially to single-trial correlations, but this was not the general tendency: most Specificity Index and Behavioural Relevance Index distributions were symmetrical. This also held for Data set 1, implying that in both data sets, any sub-set of neurons could convey the average template to approximately the same extent.

Even if there seemed to be no separate group of neurons with particularly informative responses, it is in principle possible that downstream areas only ‘listen to’ the neurons at the most informative tail of the distribution, and ignore responses from less informative neurons. To explore whether in this scenario, single-trial responses would clearly reflect the relevant cross-trial averages, we sub-sampled all recorded neurons to include only the 5 percent of each neuronal population that had emerged as most and least informative, respectively, based on the jackknifing procedure detailed above. As one would expect, the Specificity and Behavioural Relevance derived from the most informative neurons was higher than those derived from the least informative neurons. However, the difference between both was marginal, and indices for both groups of neurons fell squarely within the range of the indices we had previously obtained for the entire population. This suggests that only reading out the individually most informative neurons did not necessarily improve the reliability of the resulting population response.

In addition, by pooling stimulus pairs with large and small contrast differences into just two stimulus categories - ‘target left’ and ‘target right’ - we may have caused the resulting average templates to appear less distinctive. Specifically, difficult stimulus pairs might ‘blur the boundaries’ between average templates. To estimate to what extent the poor specificity of single-trial responses in Data Set 2 was caused by the similarity between stimulus conditions, we computed Specificity Index and Behavioural Relevance separately for difficult and easy trials. Both Specificity and Relevance increased significantly in a majority of brain areas when only taking into account stimulus pairs with large contrast differences (see Fig. S8). This suggests that in Data Set 2, average population firing rates were both specific and behaviourally relevant on a single-trial level when processing coarse stimulus information. In contrast, average population firing rates were neither specific not behaviourally relevant when animals were engaged in finer contrast discrimination. In this context, it is important to note that animals were also highly successful in discriminating these difficult stimulus pairs. This suggests that fine contrast discrimination relied on other coding modalities than average firing rates.

Another potential limiting factor of our analysis could be that in the more complex behavioural context of Data set 2, template-matching may occur in a way that could not be captured by simple correlations. As we have set out above, linear correlations are in principle the correct way to test the assumptions implied by the computation of cross-trial averages: Since an average is computed by linear operations (addition and rescaling) along one dimension, it assumes that single-trial responses vary linearly along one dimension - otherwise one would need to apply a different (non-linear and/or multi-dimensional) summary measure instead of a linear average.

However, it is still in principle possible that the linear operation of cross-trial averaging somehow reflects neuronal features of single-trial responses that are better understood in a higher-dimensional space. To explore this scenario, we repeated all previous analyses, but characterized population responses using Principal Component Analysis (PCA) via Singular Value Decomposition (SVD), and quantified their resemblance (normalized distance, see Methods) to the average template in this dimensional-reduced space. In both data sets, stimulus specificity increased marginally, but behavioural relevance decreased (Fig. S11). The decrease in behavioural relevance was particularly steep in Data set 1, indicating that raw firing rates were more instructive to the animals’ choices than dimensional-reduced features of neuronal activity (Fig. S11).

Finally, neuronal responses may reflect a multi-factorial conjunction of response preferences to a wide range of stimulus features and behavioural variables. To test this hypothesis, we quantified whether single-trial correlations to the average would become more consistent or predictive of target choice when additional variables were taken into account. As a first test, we accounted for potential modulating variables in an agnostic way by clustering the neuronal population responses from all trials according to similarity. Such clusters might reflect different spontaneously occurring processing states that the animal enters into for reasons (e.g. locomotion, satiation, learning etc.) that may remain unknown to the experimenter. Based on the Silhouette Index, which measures cluster compactness (Fig. S10), we decided to group trials into two clusters. We repeated all analyses of response specificity and behavioural relevance within each of these trial clusters. In Data set 1, the Specificity and Behavioural Relevance Indices were largely unchanged. This aligns with the fact that clusters in Data set 1 were less compact (Fig. S9A) and thus trial-grouping improved trial homogeneity less markedly. In contrast, in Data set 2, response specificity rose sharply while behavioural relevance decreased markedly (Fig. 6 B, bottom). This suggests that in this data set, accounting for potentially hidden behavioural states that might be encoded simultaneously by the same neuronal populations was not at all helpful to reveal the behavioural relevance of our stimulus-specific cross-trial averages.

As a second test, we accounted for spontaneous fluctuations of attentional state as reflected by pupil size [45, 50, 51]. To this end, we grouped the trials by pupil size and computed cross-trial averages for these sub-groups. If population responses are modulated by attentional state, examining only trials that occurred during similar attentional states should reduce unexplained variability. However, grouping trials by average pupil size did not improve specificity or behavioural relevance in Data set 2 (Fig 6 C; Data set 1 did not contain measurements of pupil size).

Together, these analyses suggest that the missing link between single-trial responses and cross-trial averages in Data set 2 is not explained by unmeasured confounding factors, non-linear interactions or lack of neurons. Rather, it appears to be a systemic feature of the data set. Fig. 7 summarizes the outcomes of different analysis approaches. In all cases, Data set 1 shows better stimulus-specificity and behavioural relevance than Data set 2. Furthermore, we show that an alternative template-matching procedure (i.e., Z-scoring the neural responses over trials, denoted by z-FRates) improves Stimulus Specificity in Data set 2, but not its Behavioural Relevance Index (and it heavily decreases Behavioural Relevance in Data set 1).

**FIG. 7.**
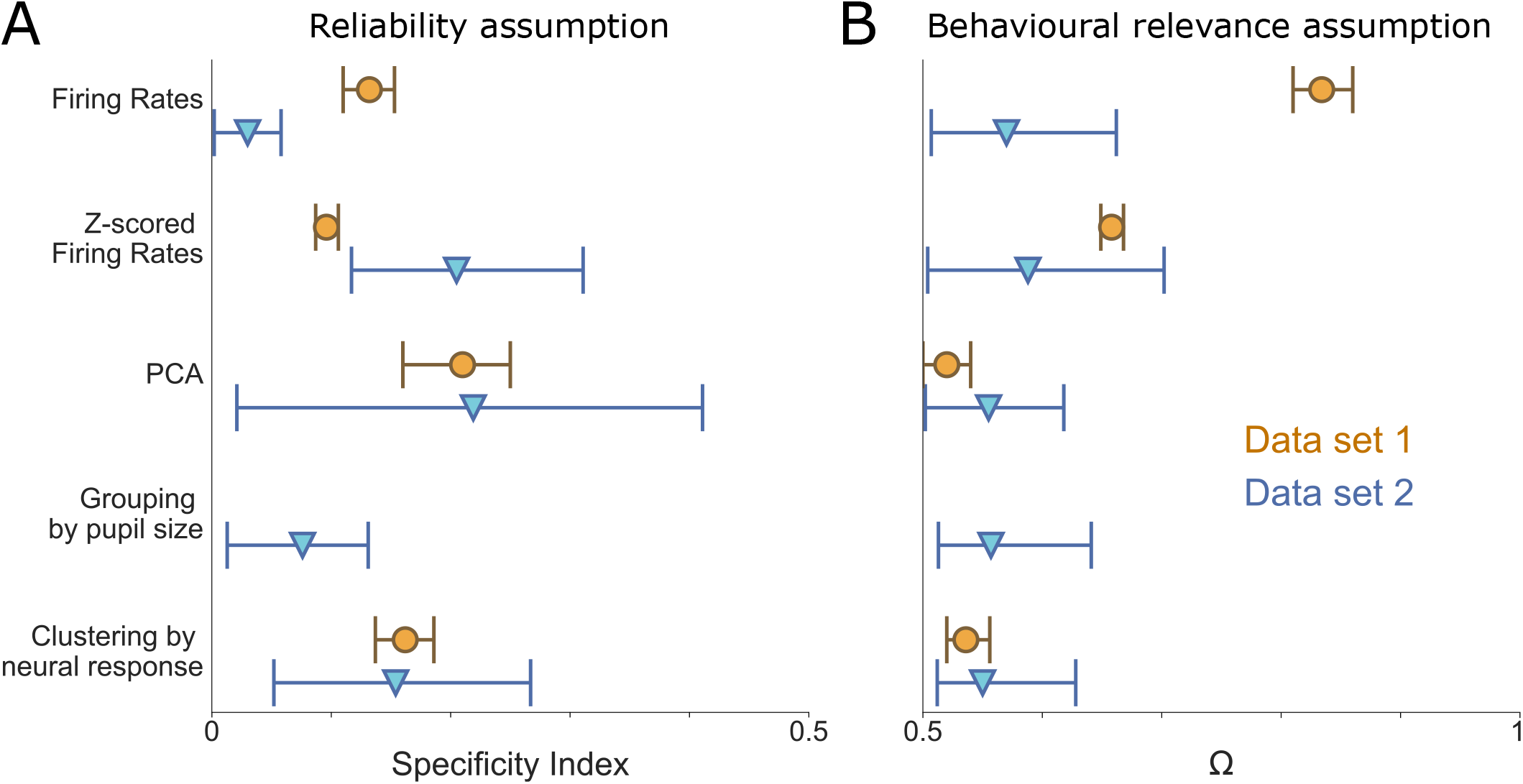
Summary of the results in both data sets.Y-axis shows the various methods used before specificity (left) and relevance (right) indices are computed.

These results raise several questions. First, what features of Data set 1 make cross-trial averages so much more representative of single-trial processing than in Data set 2? And how representative are those features compared to the range of data generated by neuroscience? To start delineating answers to these questions, we simulated how Specificity Index and Behavioural Relevance Index behave when applied to neuronal population responses with (1) different distributions of response preferences, (2) different degrees of neuronal single-trial variability and (3) different degrees of variability in the translation from neuronal responses to behavioural decision making. Specifically, we simulated a neuronal population of similar size as those recorded in both benchmark data sets (*n* = 200). For these 200 neurons, we simulated a distribution, governed by parameter *β*, of average response preferences regarding two hypothetical stimuli. When *β* = 1, response preferences were distributed completely uniformly across the spectrum from Stimulus 1 to Stimulus 2. *β <* 1 indicated a shift towards an increasingly segregated bimodal distribution, with sub-groups of neurons preferring either Stimulus 1 or 2. *β >* 1 indicated an increasingly tight unimodal Gaussian function, with all neurons responding in largely the same way regardless of the presence of Stimulus 1 or 2. Next, single-trial responses of each neuron comprised not only its ‘true’ response preference, but also varying levels of ‘noise’. Finally, based on these simulated single-trial responses, we simulated a behavioural read-out that acted as a nonlinearity, choosing the correct or incorrect behavioural output depending on how similar a given neuronal single-trial response was to the correct versus incorrect average template. This non-linearity could either act in a noise-free manner, translating single-trial responses directly to the most closely corresponding behavioural decision, or it could add some ‘decision variability’ of its own (Fig. 8a).

**FIG. 8.**
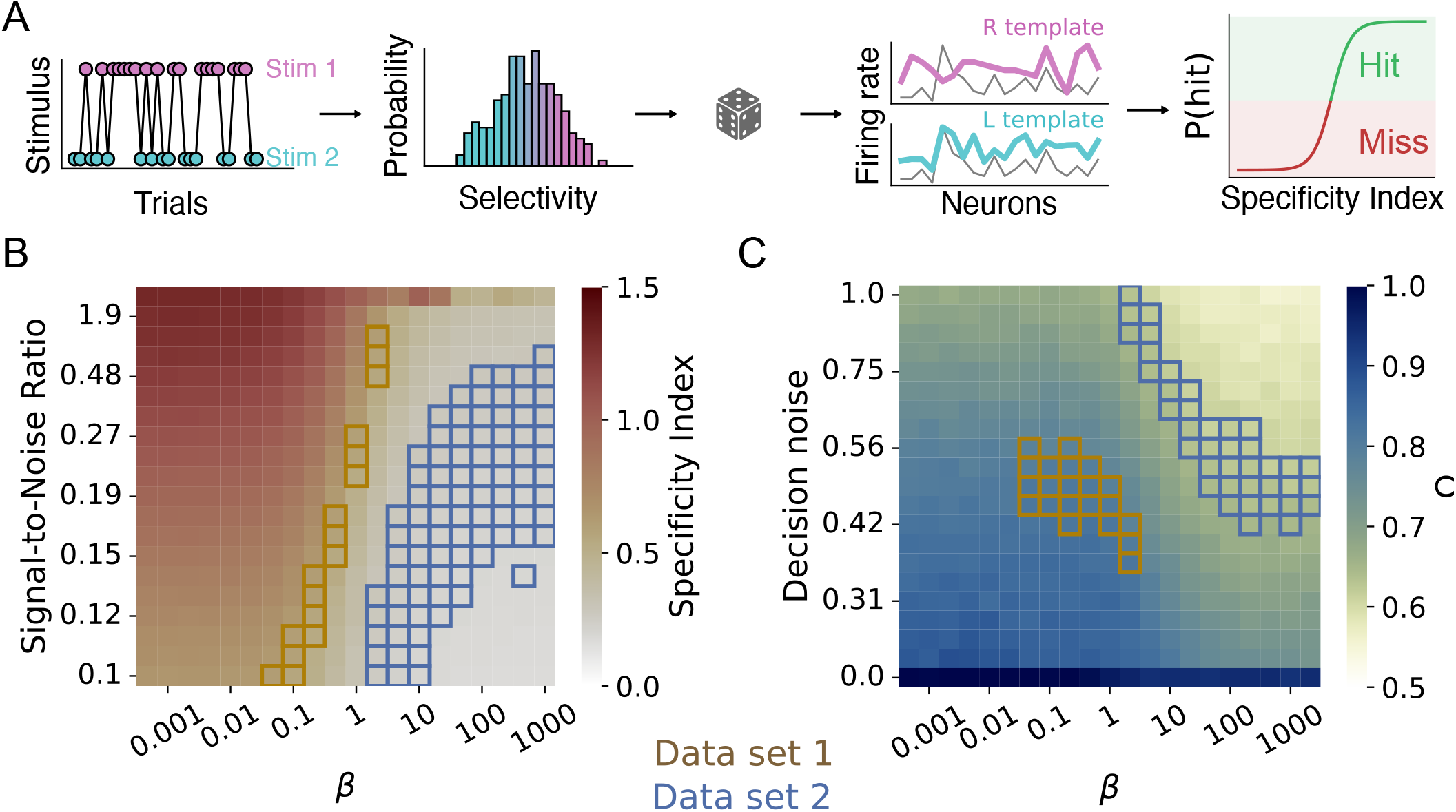
Template-matching simulation. A) Stimuli are randomly sampled from a bimodal distribution. Then, neural responses are modelled as a baseline firing rate plus a stimulus-related response (with a background noise term, parametrized by an SNR), modulated by the selectivity *β*. Finally, the choice is made by passing the Specificity Index (difference between the correlation to one stimulus minus the other) through a sigmoid with Gaussian noise. (B) Specificity Index as we vary the SNR and the selectivity (*β*) of the model neurons. We have highlighted the points in the simulation that are compatible with the experimental data sets (color coded as indicated in the legend). (C) Same as B), but for the Behavioral Relevance Index (Ω). In this case, we varied the noise intensity of the decision-making process, for a fixed intermediate SNR.

These simulations showed that the Specificity Index will rise steeply when the distribution of neuronal response preferences is reasonably spread out, e.g. in the form of a uniform distribution (*β* = 1) or a bimodal distribution (*β*

¡ 1). In contrast, the Specificity Index will remain low as soon as neuronal response preferences are more stereotyped than stimulus-specific (*β* ¿ 1). Interestingly, this overall pattern was only marginally dependent on the amount of single-trial ‘noise’ added to the simulated average response preferences (Fig. 8B). This suggests that the main factor determining the Specificity of single-trial responses is a broad coverage of different response preferences by the studied neuronal population. Comparing the median Specificity Index of Data set 1 and 2 to these simulations (Fig. 8B) suggests that neuronal responses in Data Set 1 should distributed at least somewhat bimodally between its two stimulus conditions, while those in Data Set 2 should be less distinguishable from each other. The real distributions of neuronal response preferences in both data sets confirmed these predictions (Fig. S13). These simulations therefore suggest that the better Specificity Index of cross-trial averages in Data set 1 than in Data set 2 largely hinges on the broader distribution of neuronal response preferences.

We next explored how our Behavioural Relevance Index reflected the interplay between the distribution of neuronal response preferences, and the variability in the decision making process itself. Like the Specificity Index, Behavioural Relevance increased with a broader distribution of response preferences, as well as (unsurprisingly), with smaller variability of decision making (Fig. 8C). Based on their median Behavioural Relevance Indices, our two benchmark data sets would appear to occupy distinct areas of the parameter landscape. The Behavioural Relevance in Data set 1 suggests that it operates in a regime with an, at least somewhat, bimodal distribution of neuronal preferences and a moderate variability of behavioural decision making, meaning that single-trial correlations to the average template predict behavioural decisions quite well. In contrast, Data set 2 might feature either a tight unimodal distribution of neuronal response preferences, combined with similarly faithful decision making as Data set 1 - or a more spreadout, uniform distribution of neuronal preferences, but with extremely noisy decision making. Given the response distributions shown in (Fig. S13), we assume that both the clustering of neuronal response preferences and genuine decision-making variability contribute to the low Behavioural Relevance observed in Data set 2.

Together, these simulations demonstrate three main outcomes. First, the Specificity and Behavioural Relevance Index are able to correctly pick up the neuro-behavioural features they were designed to reflect - stimulus-specific neuronal response profiles in the case of the Specificity Index, and consistent decision criteria based on these neuronal response profiles in the case of the Behavioural Relevance Index. Second, our two benchmark data sets occupy distinct and complementary spaces within the parameter landscape of neuronal and behavioural variability, with the lower effectiveness of cross-trial averages in Data Set 2 being most likely due to a tighter distribution of neuronal preferences and higher variability of decision making. Finally, based on the Specificity and Behavioural Relevance indices computed in real data, simulations like the one presented here can in fact be used to hypothesize about the neuronal and behavioural mechanisms that boost or bust the applicability of cross-trial averages to single-trial responses in a specific experiment. For instance, our simulation highlighted the broad distribution of neuronal response preferences, rather than the magnitude of single-trial variability, as the main factor that makes cross-trial averages more meaningful in Data set 1 than in Data set 2.

## Discussion

The present study set out to formulate an explicit statistical test to determine how reflective average population responses are of neuronal processing in different contexts. To this end, it posits two testable assumptions that should hold if cross-trial averages are computationally relevant:

1. single-trial responses should be sufficiently reliable and specific to resemble the correct average template (if average templates are not reflected in single-trial responses consistently, they cannot reflect computationally relevant aspects of single-trial responses)

2. better-matching single-trial responses should evoke more efficient behaviour (if average templates represent the true neuronal signal, then single-trial responses that better resemble them should by definition provide a better signal-to-noise ratio).

To directly quantify to what extent these two conditions are fulfilled in a given data set, we introduced two simple metrics: (1) The Specificity Index, which captures whether a single-trial response is more related to the average response of the true experimental condition than to the average responses of other experimental conditions; and (2) The Behavioural Relevance Index, which reflects whether higher single-trial correlations to the correct average template result in more behaviourally successful trials.

With this test, we aim to bridge an in our view crucial gap in the way neuroscience is currently practiced: The disconnect between our explicit knowledge that neuronal population activity typically evolves continuously according to non-linear dynamics, which are often poorly captured by cross-trial averaging ([25, 52–55]), and the implicit assumptions we accept when nevertheless treating cross-trial averages of neuronal activity as an informative summary metric. Our two metrics are easy to compute and thereby allow researchers to explicitly determine whether their trial-average metrics are likely to be sufficiently reliable and behaviourally relevant to play a functional role in a given experimental context.

To establish the general applicability of our metrics, we benchmarked them on two complementary data sets, featuring tight experimental control of neuronal stimulation and highly trained, simple behavioural responses in the first data set, and less tightly controlled visual stimulation with a more variable and naturalistic behavioural output in the second data set. In these benchmark tests, we find that the two assumptions of cross-trial averaging are largely fulfilled in only the first data set. This success is surprising because optogenetic stimulation targeted a somewhat overlapping but randomly selected population of 5 to 150 neurons in each new trial. Thus, even though optogenetic stimulation varied randomly, it seemingly managed to recruit a reproducible network of neurons within the analysis time window of 500 ms post-stimulation. This suggests that the population responses highlighted by our analyses of Data set 1 rely on ‘hub neurons’ that are activated by various different stimulation patterns. Moreover, this is in line with the encoding of stimulus responses across trials by a consistently weighted set of neurons [34].

In contrast, in Data set 2, single-trial correlations were robust but not stimulus-specific, implying that they mostly reflected factors such as baseline firing rates rather than conveying stimulus information. Correcting for such baseline differences in firing rate, as done by PCA, did not improve the Specificity index of single-trial correlations. Moreover, single-trial responses that better resembled the correct template hardly increased an animal’s chance of choosing the correct target. Further analyses indicated that these results would not improve with more neurons, or by taking into account complementary variables such as behavioural state. This suggests that in Data set 2, average population responses were not the central mechanism driving perceptual decision-making.

The disparity of outcomes between the two data sets examined here could be due to several factors. In many ways, Data set 1 offers an ideal case for average population responses to play a functional role: Stimulation is highly controlled, and takes place directly within the recorded brain area rather than being relayed across several synapses; behavioural responses (in the form of licking) are short and stereotyped, reducing movement-related neuronal dynamics; and animals are explicitly trained to detect a difference in the average amount of neuronal activity within S1. In other words, even if sensory stimuli were typically not encoded in average S1 population firing rates, animals in Data set 1 may have essentially learned to ‘count S1 spikes for reward’.

By comparison, Data set 2 features less controlled visual stimulation since animals can look at either of the two stimuli freely; modulation of the stimulus signal by several synaptic relays; additional neuronal dynamics driven by increased and more complex spontaneous movement in the form of wheel turning; and of course the fact that animals are free to process the difference in grating contrast in ways other than average population firing. Thus, while the two data sets examined here are clearly not sufficient to draw general conclusions, our results may point towards a scenario where cross-trial averages are more computationally relevant in settings featuring strong stimulus control and expert or over-trained behaviours. This would mean that especially in order to understand more naturalistic neuronal computations, cross-trial averages might not be helpful.

An alternative possibility is that we underestimated the computational role of cross-trial averages in Data set 2 due to idiosyncrasies of the paradigm and of our analyses. First, task-relevant stimulus information in Data set 2 had to be computed by comparing visual inputs across brain hemispheres, but we only had access to neuronal activity from one hemisphere. Thus, recording from both hemispheres might have yielded more informative population templates. Nevertheless, the animal is proficient in this visual 2AFC task, which means that even if some brain areas (e.g. primary visual cortex) do not integrate information from both hemi-fields, some downstream area(s) should receive the result of the cross-hemisphere computations needed to initiate the correct behavioural response. Since this data set is arguably the most complete set of neuronal recordings to date regarding the number of recorded brain areas, it seems unlikely that not a single area consistently represents integrated stimulus information from both hemi-fields.

A popular notion is that specific features of neuronal processing may be encoded by ‘super-groups’ of neurons that are dedicated most reliably to the feature in question. While such a segregation of neuronal sub-populations has previously been successful [48], our control analyses suggested that not only were there no clearly identifiable ‘super-coder’ neurons, but selecting only for the most informative neurons also brought no significant benefit in terms of Specificity and Behavioural Relevance. This suggests that at least in this context, reading out a more eclectic mix of more and less stimulus-specific neuronal responses would actually benefit information decoding.

Finally, the stimulus-related response templates explored here may generally underestimate the computational power of average responses by ignoring the many stimulus and behavioural factors at play at any moment in time [5, 16, 17, 43, 45–47, 56], only some of which will be known or accessible to the experimenter. This can make neuronal responses appear highly unpredictable, while they are actually shaped systematically and reliably by a set of unmeasured, or ‘latent’, variables.

We investigated this idea in two different ways. First, we grouped trials by pupil size, which is known to reflect spontaneous fluctuations in attentional state that strongly shape neuronal population activity [45, 50, 51]. Second, to account for modulating variables in a more agnostic way, we searched for distinct trial clusters that featured similar neuronal population responses. Such clusters might reflect different processing states that the animal enters into for reasons (e.g. locomotion, satiation, learning etc.) that may remain unknown to the experimenter. If either of these variables formed part of the ‘multi-factorial average response curve’ of the recorded neurons, then only considering trials recorded during a similar attentional state (as measured by pupil size) or within the same trial cluster should increase the specificity and behavioural relevance of the resulting cross-trial averages by removing one source of response modulation that was previously unaccounted for. This was the case in Data set 1 but not in Data set 2, suggesting that even when more latent variables are accounted for, averages may still not be the most accurate way to reflect information processed by the brain.

In addition, several recent papers have argued that factors such as stimulus properties, behavioural choices, and retrieved memories are encoded along largely orthogonal dimensions in neuronal response space [8, 57, 58]. If this is true, then extracting cross-trial averages via a dimensionality-reduction technique like PCA should enable us to dissociate them from other, orthogonally-coded, types of information, and thereby significantly improve their computational relevance even in the presence of other modulating factors. This scenario held to some extent for Data set 1, where single-trial response vectors as extracted by PCA became more specific to the average template, but not more behaviourally relevant. In Data set 2, applying PCA did not result in any substantial improvements. It is possible that these outcomes depend on the choice of dimensionality-reduction technique. We chose PCA as a benchmark due to its simplicity and ubiquitous use, but other approaches like non-Negative Matrix Factorization

[59] might yield different results [60]. If they were to prove more successful, this would argue in favour of analyses characterizing average neuronal response preferences simultaneously for multiple, potentially non-linearly interacting factors [61]. In this case, we would suggest that the neuroscience community abandons single-feature response averages in favour of average multi-feature response ‘landscapes’. This would involve finding routine metrics to track ubiquitous latent variables like behavioural state [45, 47, 62–64] throughout a wide range of experiments.

Crucially, our results suggest that the utility of trial-averaged responses can vary dramatically across different contexts. The relevance of trial-averaging is likely to shift depending on behavioural context, stimuli, species – as well as the aspect of neuronal activity that is averaged, such as neuronal firing rates, firing phase, coherence etc. Similarly, dimensionality-reduction techniques may improve the reliability of cross-trial averages in some cases (such as Data set 1), but not in others (as in Data set 2).

[31]. This notion was also supported by our simulations exploring how Specificity and Behavioural Relevance might scale across data that contained variable amounts of neuronal and behavioural variability as well as different distributions of neuronal response preferences. Based on these simulations, we predict that, rather than neuronal or even behavioural choice variability, the sheer diversity of neuronal response preferences is the most crucial factor rendering cross-trial averages relevant to single-trial computations in a given context - and meaningless in another. We aim to test this prediction more formally once Specificity and Behavioural Relevance are established for a hopefully fast-growing number of neuronal data sets.

Most importantly, we encourage researchers to compute simple ‘rule-of-thumb’ metrics such as the Specificity Index and Behavioural Relevance Index in order to estimate what computational role cross-trial averages play in the experimental paradigms, neuronal computations and neuronal response metrics that they study. Over time, we hope that this practice will generate a ‘map’ of contexts in which cross-trial averages are computationally meaningful - and motivate the field to restrict the computation of cross-trial averages to cases when they are in fact relevant to the brain.

If classical trial-averaged population responses appear largely irrelevant to ongoing neuronal computations at least in some contexts, how then could stimulus and target choice information be encoded in such cases? First, information may be encoded mostly in joint neuronal dynamics that are not captured by static (single- or multi-feature) response preferences. Analyses that take into account such dynamics, e.g. by tracking and/or tolerating ongoing rotations and translations in neuronal space [64–67] or by explicitly including shared variability in their readout [68, 69], often provide vastly more informative and stable neuronal representations [65, 66]. Consistent with this, the decoder analyses (Fig. 4) extracted information more successfully – most likely because decoders rely on co-variability and co-dependencies between input data and the class labels, which are smoothed over by trial-averaging.

Second, while here we have tested cross-trial averages of population firing rates as a benchmark of basic analysis practices in neuroscience, other aspects of neuronal activity might be more informative - and as a result also potentially lead to more informative cross-trial averages. For instance, transiently emerging functional assemblies [70–72], phase relationships between neuronal sub-populations [73–75] or the relative timing of action potentials [53, 76, 77] may provide an avenue of information transmission that is entirely complementary to population firing rates.

Finally, by tracking more stimulus and behavioural variables at the same time, we can further explore how they dynamically influence overlapping and separate aspects of neuronal activity ([9, 62, 63, 78–81]). Over time, we hope that this will shape our understanding of neuronal activity as an ongoing interaction rather than a static snapshot. No matter which of these approaches turns out to be most successful, it is important to recognize that time-averaged population responses may, at least in some contexts, not be a fitting way to describe how the brain represents information.

## Acknowledgements

We thank Jonathan Pillow, Viola Priesemann, Nick Steinmetz, Cyrille Rossant, Miles Wells, Joshua Gold and Mike X Cohen for valuable input on earlier versions of the manuscript.

## SUPPLEMENTARY MATERIAL

## METHODS

We have released all the scripts and data files to reproduce these analyses, they can be found at the following URL: https://github.com/atlaie/BrainAveraging. They are written in Python 3 and rely on several libraries.

### Specificity index

With the intent of characterizing whether the neural response is more similar to the appropriate template (i.e., the one corresponding to the stimulus that was actually presented in that trial) or the other one, we introduced a simple quantity we termed specificity index. It is defined as:

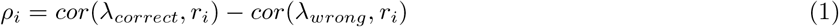

where *cor* is the Pearson correlation, *λ* denotes a given neural template and *r*_*i*_ is the population vector of the *i*^*th*^ trial. Thus, the specificity index captures the differential similarity of a given neural response to each of the templates. It is key to note that, given that the Pearson correlation is bounded between −1 and 1, the specificity index can attain values between −2 and 2 and, as we were just interested in its sign and global tendencies, we did not introduce any normalization factor.

### Behavioral relevance

As a way to quantify the overlap between the hit and miss distributions we basically used Vargha-Delaney’s *A* effect size [36] (also known as *measure of stochastic superiority*). This is an effect size derived from the Mann-Whitney U-test

–a non-parametric statistical test that is particularly useful when distributions are not Gaussian [82]. Furthermore, *A* is especially interpretable. As it is related to the *U* statistic, it can be thought of [83] as the probability of a randomly selected point from one distribution (*X*) being higher than another randomly selected point from the other distribution (*Y*). Before computing the *U* -statistic, we have to:

1. Pool all data points into one group and sort them from low to high values.

2. Assign ranks to each sorted data point. If there is a tie (i.e., two repeated values), their rank is taken to be the average of the ranks for the entire pooled group.

3. Compute *R*_*X*_ and *R*_*Y*_ as the sum of the ranks of each of the groups. Finally, the *U* −statistic will be given by:

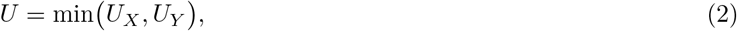

with 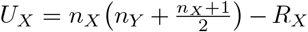 and equivalently if we flip the *X* and *Y* labels. Having computed the U-statistic, our measure of Behavioral Relevance (Ω) is given by

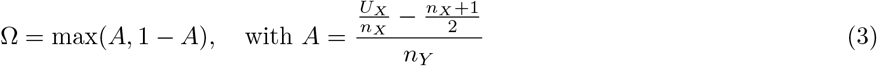

Thus, Ω is bounded between 0.5 and 1. If there is no overlap, Ω = 1. In this extreme case, one distribution would have complete stochastic dominance over the other. If *X* and *Y* are totally overlapping, Ω = 0.5 and, thus, the more its value deviates from 0.5, the less overlapping the distributions are. Note that we dropped the subscripts for Ω; this is due to its definition being symmetrical for *X* and *Y*, because of the *max*(·) operation. This can be easily shown, as:

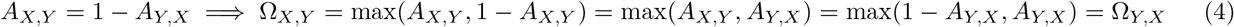

Thus, the definition of the Behavioral Relevance index is independent of the order in which we take the statistical test.

### Yule-Kendall index

In order to assess how symmetrical the jackknifed distributions shown in Fig. S6 are, we relied on the Yule-Kendall index, which is computed as:

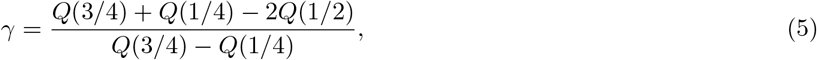

where *Q* is the quantile function. We chose this measure because it works for non-normal distributions and because it is non-dimensional (thus allowing direct comparison between data sets).

### Decoders

In order to select the most informative brain areas for Data set 2 in a data-driven way, we made use of different decoders. For each experimental session, there are several recorded regions. Thus, we trained independent decoders using the single-trial population vector for each region. The labels to be predicted would be either choice (left wheel turn, right wheel turn or no movement) or stimulus (right-higher contrast, left-higher contrast, both equal).

We split the data following a 80 − 20 ratio (train-test). Given the imbalanced nature of the dataset, we used a stratified (i.e., ensuring that all classes are fairly represented across each fold) 10−repeated 5−fold Cross-Validation to fine-tune each model’s hyperparameters using a Bayesian approach and checked that the model performance was above majority class (i.e., always predicting the most abundant label) and random models.

In the following, let assume we want to classify *N* input variables (*x*_*i*_, with *i* = 1, …, *N*) into target variables that can belong to *K* classes (i.e., *y*_*i*_ ∈ {1, …, *K*}).

#### Bayesian optimization

Hyperparameter optimization is arguably one of the bottlenecks when deploying Machine Learning techniques. Fortunately, in the last years, the field is rapidly evolving and we now have access to quick and intuitive libraries that alleviate greatly these tasks. One of these libraries is *Optuna* [84]. We relied on Tree-structured Parzen Estimators (TPEs) [85] to speed up hyperparameter search.

For each model, we chose different hyperparameters to be optimized, which we will detail in the following descriptions. We relied on the SciKit-Learn package in Python [86] to implement all of the classifiers.

#### Gaussian Naive Bayes

Naive Bayes assumes independent features [87]. In terms of the covariance matrices, this model assumes they are diagonal. Particularly, the Gaussian Naive Bayes (GNB) model assumes that the class-conditional densities are normally distributed as:

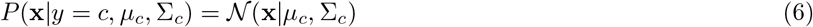

where *μ* and Σ are the class-specific mean vector and class-specific covariance matrix, respectively. In order to compute the class posterior, we can simply make use of Bayes’ theorem

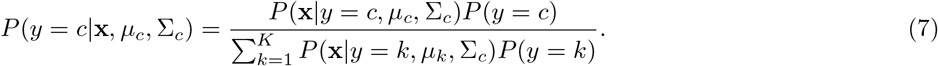

So, in order to classify **x** into a class *c*:

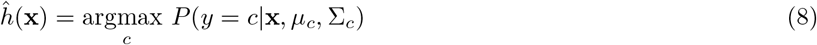

For this classifier, we did not perform any hyperparameter tuning.

#### k-Nearest Neighbors

This model relies on the assumption that the closer two points are, the more similar they are and the more likely it is that they belong to the same class. It is implemented as follows: for any given point *x*_*i*_, compute its distance to the rest of the points; then, select the *k* points that are closest; out of those, apply a majority vote rule. As a summary: the most prevalent class of the *k* closest points to *x*_*i*_ will determine the class that we predict for it.

For this classifier, we optimized the number of nearest neighbors and the leaf size of the k-d Tree algorithm.

#### Random Forest

This model makes use of several Decision Trees (DTs) to solve supervised learning problems. DTs are non-parametric models that split the data using a given number (depth) of conditional steps - populated with at least with some number of data points (minimum sample split). When we aggregate several DTs (using a technique known as bootstrap aggregating or *bagging*), we end up with a *Random Forest* (RF). Specifically, we sample with replacement from the data and feed these subsamples to different trees. Finally, we use a majority vote for each tree’s output as the RF prediction. This procedure reduces the variance in the model (i.e., mitigates overfitting) but has a minimal effect on its bias (i.e., underfitting is still a risk). Therefore, ideally we would like to use DTs that are unstable (high variance) but very little bias. In order to check the model performance, we compute its out-of-bag (oob) error - cases in the training data that are not in a significant part of the bootstrapped samples.

For this model, we optimized the number of DTs, their maximum depth and the minimum sample split.

#### Generalized Linear Model

We trained a multinomial Generalized Linear Model (GLM) with a L2-regularization. This model states that the probability of a particular data point *y*_*i*_ belonging to class *c* is given by:

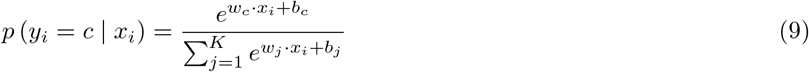

After having the probabilities of *y*_*i*_ belonging to each class *c*, the highest one will be taken to be 1 and the rest will be set to 0. Therefore, the objective is to find the weight vector *w*_*c*_ that minimizes the distance between the predicted (*ŷ* _*i*_) and the actual (*y*_*i*_) class labels by optimizing (in this case, minimizing) the following loss function:

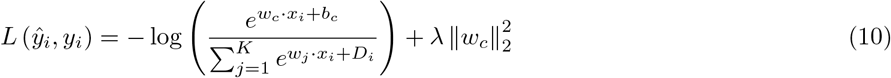

where 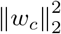 is the L2-norm of the weight vector for class *c*, accounting for the L2-regularization term, with *λ* modulating its strength. The inverse of *λ* (*C* = 1*/λ*) is the hyperparameter we optimized for this model.

#### Support Vector Machines

This class of models rely on finding the hypersurface that maximizes the separation between classes in the data. To this end, it is important to first find the *support vectors* (SVs), which are the closest ones to the surface. There are two main types of Support Vector Machines (SVMs): linear SVMs and kernel-based SVMs. The difference is that, in the former, the separating hypersurface is a hyperplane (i.e., a non-curved hypersurface), while, in the latter, it is is allowed to be curved. Mathematically, the *linear* version of this model is the solution to the following optimization problem:

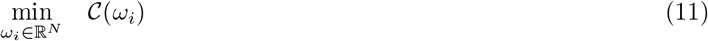

where

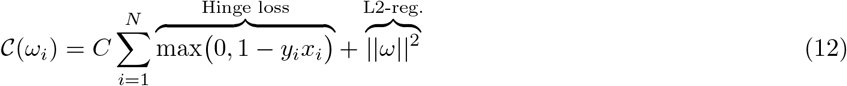

In order to arrive to the equivalent one for the non-linear SVM, we can take advantage of the fact that this problem satisfies some conditions [88] that allow us to construct its dual form. It takes the form of:

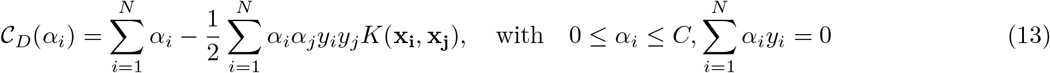

where *K*(**x**_**i**_, **x**_**j**_) is the kernel (it can be linear or not) and *C* is the regularization strength. In this work, we chose Radial Basis Functions (RBF) as our kernel. These are given by:

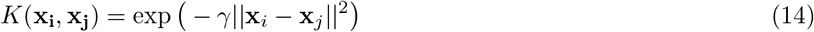

where *γ* is the reach parameter (i.e., how far we want two separate points influence each other).

For these decoders, we optimized the regularization strength (*C*) and, only for the *RBF*, the reach coefficient (*γ*).

### Mutual Information

After having trained each decoder, we separately computed the Mutual Information between the predicted and the test class labels, as a proxy of the amount of stimulus – or choice – information there was in the population vector.

This quantity is defined in the context of classical Information Theory [89, 90] and we can compute it for two discrete stochastic variables *X* and *Y* . Assuming these have a joint probability mass function given by *p*_*X,Y*_ (*x, y*) = *P* (*Y* = *y* | *X* = *x*) · *P* (*X* = *x*) and that each of them follows a marginal probability distribution given by *p*_*X*_ = *y*∈*Y p*_*X,Y*_ (*x, y*), one can mathematically define the Mutual Information between *X* and *Y* as:

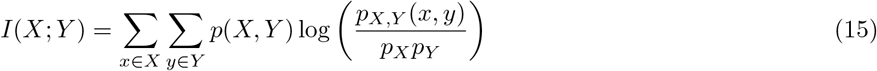

Intuitively, one can understand *I*(*X*; *Y*) as the uncertainty reduction in *X* that follows if *Y* is measured (or vice versa, as *I*(*X*; *Y*) is invariant when swapping *X* and *Y*). If (and only if) they are independent of each other, then *I*(*X*; *Y*) = 0. Therefore, this is a strictly non-negative quantity. It is noteworthy that *I*(*X*; *Y*) captures all linear and nonlinear dependencies between *X* and *Y*, thus generalizing the notion of correlation measures. For further discussion of this measure, see [91, 92].

### Hierarchical clustering

After having computed the resulting Mutual Information for all areas, sessions and decoders, we aimed to check the stability of the selection of the most informative brain areas, so that results were not highly dependent on which model we used to choose them. To do that, we used an unsupervised method, known as hierarchical clustering. Particularly, our vectors were the aggregated information (stimulus + choice) that each decoder extracted, over all brain areas. Once we have the pairwise distance between points (proximity matrix), this method can be understood in an iterative manner: merge the closest points in a cluster, then merge the closest clusters and repeat until only a single cluster (encompassing all points) remains. For Fig. 4, we used the Euclidean metric to compute the proximity matrix.

To implement this algorithm, we relied on the Seaborn [93] (Python package) implementation of *clustermap*.

### Elbow method

In order to select a threshold when selecting the task-related areas based on their stimulus and choice information (Figure 4 in the main text), we used the data to compute the Kernel Density Estimate, via Gaussian kernels [94]. After having extracted these, we used the method discussed in [95] to find the point of maximum curvature. We made use of the kneed Python package, implemented by the same authors [95].

### Surrogate models

#### Calcium imaging data

For this data set, we pooled together those trials within the same stimulus set: on the one hand, when stimulation was given; on the other hand, those trials without any stimulus. For each of those trial groups, and for each neuron, we built a Gaussian distribution with mean and variance given by the trial-average and trial-variance. We then sampled 200 random values for each stimulus set and correlated each of them with the template.

#### Spike data

We were interested in comparing the experimental neuronal population response with a downsampled version of the trial-averaged template. To do that, we built our surrogate models by constructing *N* (= 100) random vector with the following constraints:

1. Its size is equal to the number of neurons comprising the neural population for that area and that session.

2. The probability that at n spikes are allocated at a particular location m (i.e., that neuron m has spiked n times) is given by 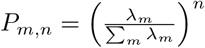, where *λ*_*m*_ is the *m*^*th*^ element of the template vector.

3. The total number of spikes is constant and equal to the total recorded number of spikes for that area and that session.

By imposing these constraints, we are testing the alternative hypothesis that neurons are independent from each other (uncorrelated) and it is therefore equivalent to keeping the single-trial population statistical response, while scrambling across trials. This is also the same as drawing single-neuron responses from the underlying template distribution following a Poisson process. Thus, the bootstrapped responses contain the same number of spikes as the original trial, but the neurons that produced these spikes are randomly chosen according to their probability of occurrence in the average template.

### Templates and distances in PCA space

As an alternative to Pearson’s correlation, we applied Principal Component Analysis (PCA) [96]. We chose PCA over non-Negative Matrix Factorization [97] or other more advanced dimensionality reduction techniques such as LFADS [68] or PSID [64] because we wanted to keep all analyses as general as we possibly could. Thus, we computed the truncated Singular Value Decomposition (tSVD) [98] for the matrix consisting of Z-scored single-trial population vectors, for a given area and session. Then, we extracted the knee (elbow) using the aforementioned method, to select the number of components based on the variance explained. After the number of components has been selected, we projected each single-trial into this (dimensionally-reduced) space and computed the Euclidean distance between this new vector and the template (also projected into this space). We normalized by the distance between the projection of the two templates in this new space.

### Specificity Index for PCA distances

Since in PCA analyses we dealt with distances rather than correlations (i.e., differences rather than similarities), we inverted the computation of the Specificity Index in this context so that positive values continued to signify a stronger relation between single trial response and correct template than incorrect template. The corresponding formula is:

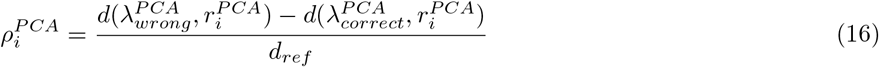

where *d* stands for Euclidean distance and 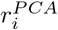 is the PCA-projected version of the population vector measured in the *i*^*th*^ trial; 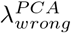 and 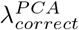 are the PCA-projected version of the trial-average templates (wrong and correct, respectively). Finally, 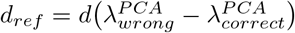, which we took to be the reference distance.

### Pupil size

As a way to account for the animal’s behavioral state, we grouped together trials that had a similar pupil size. Firstly, we low-pass filtered the recorded pupil size (that came as an output of DeepLabCut [99]) using a Butterworth filter [100] of order 4. We chose to filter out any frequency above 1*Hz*, as we were interested in slow variations in pupil size, which have been linked to attentional state [101, 102]. We then computed the mean pupil size in the same time window we used for the neural analyses (200 *ms* after stimulus presentation). Finally, we ranked all sizes over trials and grouped them in brackets of 10 percentiles (e.g., trials with a pupil size between the 32^*th*^ percentile and the 42^*th*^ one would be grouped). Within the selected group, we repeated the template-matching algorithm that we used in the main text, tailoring the average to those trials that shared a similar pupil size.

### k-Means clustering

In order to group trials according to similarity in the recorded neural response, we used *k*-Means clustering. This algorithm, after randomly initializing *k* centroids (one per cluster), is as follows: (1) Compute the distance between each data point and these *k* centroids. (2) Each point will belong to the cluster with the closest centroid. (3) New centroids will be given by the actual points belonging to a cluster.(3) Repeat until convergence (when centroids move no more). As this is an unsupervised method, we used the elbow method to select the number of *k* centroids in which to cluster the data S7. Finally, as we did for the pupil size grouping, we repeated the template-matching algorithm that we used in the main text, tailoring the average to those trials that belonged to the same cluster.

### Ideal template-matching simulation

In order to systematically explore the two assumptions that we tested in both experimental data sets, we implemented a simple model. We begin by generating two stimuli, denoted as *S*_1_ and *S*_2_. We assumed, for simplicity, that they can only attain two possible values (0 and 1). For each of the stimuli, a series of trials are generated. Each trial is an independent and identically distributed draw from the Bernoulli distribution. Thus, for each stimulus *S* with parameter *p*, the probability mass function is:

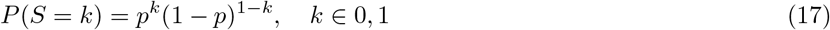

Where: *P* (*S* = *k*) is the probability that *S* takes the value *k, p* is the probability of drawing that value and (1 − *p*) is the probability of drawing the other one.

In this simulation, we chose *p*_1_ = *p*_2_ = 0.5, resulting in two possible equally likely stimuli.

Then, we simulated the activity of a population of neurons in response to different stimuli. We assumed that each neuron *j* has a certain baseline firing rate 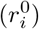 and a certain stimulus gain 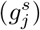, both of them sampled from the normal distribution. Both distributions, are given by N (5, 1). Moreover, each neuron has an intrinsic selectivity value (*ξ*_*j*_), which determines how the neuron’s firing rate changes depending on the difference between the two stimuli. The selectivity values were generated from a Beta distribution, controlled by the parameter *β*.

Specifically, let *ξ* denote the vector of selectivities all neurons, with *ξ*_*i*_ indicating the selectivity of neuron *i*. Then, each *ξ*_*i*_ is generated as follows:

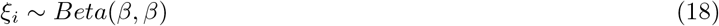

The selectivity values are then shifted and scaled to lie within [−1, 1].

For each trial, the firing rate of each neuron depends on its selectivity and the difference between the stimuli. The difference between the stimuli (Δ*S*), is computed for each trial, resulting in a sequence of stimulus differences.

The firing rate for each neuron *j* on trial *i* is then calculated as follows:

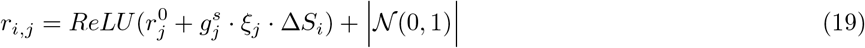

where 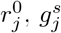 and *ξ*_*j*_ are the baseline firing rate and stimulus gain the selectivity for neuron *j*; Δ*S*_*i*_ is the stimulus difference in trial *i*. Then, we pass the computed firing rate through a ReLU in order to add a simple non-linearity and to ensure positivity.

Finally, the firing rates are used to generate spikes following a Poisson distribution. For each neuron *j* on a trial *i*, the number of spikes, denoted as *s*_*i,j*_, is generated as:

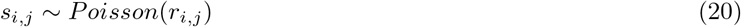

After we had the population response to the presented stimulus, we proceeded as before: we computed a “left” and “right” template (i.e., Δ*S* = 1 or Δ*S* = −1); then, for a single-trial, we computed the similarity of the population vector in that trial with each of the templates (“correct” and “wrong”). The difference in between these is the Specificity Index.

In order to model the decision-making process, we assumed that the relevant quantity to make a decision was the Specificity Index, as it follows from the template-matching procedure. We assumed there was a noisy process on top of the template-matching. Then, for a given trial, the noisy Specificity Index (*Ψ*^′^) is simply:

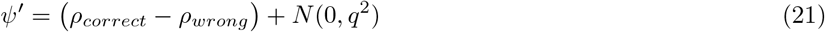

Then, and the corresponding outcome is given by:

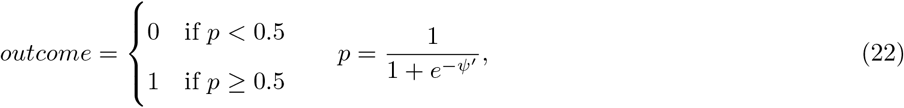

Thus, if the probability value is greater than 0.5, that trial is considered a hit (1); otherwise, it is a miss (0).

Crucially, we added an stochastic component into the decision-making, which means that even for trials with the same Specificity Index, different decisions can be made.

## SUPPLEMENTARY RESULTS

### Figure 5

To estimate stimulus-specificity, we computed single-trial correlations to the incorrect template. These resulting correlations are marginally lower than those for the correct template (Fig. 5A; difference between median correlations: 0.03 ± 0.02 across areas; t-test for dependent samples; *n* = 89 to 3560 trials per area; *t* = 3.0 to 27.2; all *p <* 0.01, corrected for multiple comparisons using FDR with a family-wise error rate of *α* = 0.05). However, the correlation difference (0.03 ± 0.02 across areas) is so small compared to the spread of single-trial correlations (standard deviation: 0.04 to 0.39) that correlations to the correct and incorrect templates were largely indistinguishable. Furthermore, task-relevant brain areas did not show more specific correlations than control areas (mean correlation difference for task-relevant areas: 0.031; *n* = 9; for comparison areas: 0.022; *n* = 62; Welch’s t-test: *t* = 1.17, *p* = 0.27).Thus, correlations between single-trial responses and template are largely stimulus-independent. To quantify this, we defined the specificity index, which measures the single-trial correlation to the correct template minus the incorrect template (Fig. 5B). Most values are positive, indicating that single-trial responses were generally more similar to the correct than incorrect template (t-test for difference from zero; *n* = 89 to 3560 trials per cortical area; *t* = 3.0*to*27.2; all *p <* 0.01, corrected for multiple comparisons). However, the distributions rarely exceeded 0.1 (Fig. 5B). Therefore, across the examined brain areas, single-trial responses were barely more similar to the correct template than to the incorrect one.

To test if the match between single-trial responses and the correct template predicted target choices, we split by hit (trials where the animal chose the correct target) and miss trials (no response or wrong target). Single-trial correlations were lower in miss trials than in hit trials across most brain areas, suggesting that a better match to the average template did indeed tend to produce hit trials more often (Fig. 5C). However, overall the difference between the correlations was small, as quantified by the Behavioral Relevance Ω (Fig. 5E). There were some exceptions (MOs, Ω = 0.66, *p* = 0.0142, *n*_*hit*_ = 2057, *n*_*miss*_ = 667), potentially because hit and miss trials are associated with fundamentally different motor responses (miss trials include trials with no choice). Yet the overall pattern suggests that template matching has low - and inconsistent - predictive power regarding perceptual decision-making. It is however possible that the important factor for perceptual decision making is not the overall correlation between the single-trial response and the correct response template, but whether it resembles the correct more than the incorrect template. However, splitting specificity indices by hit and miss trials again shows no consistent difference (Fig. 5D), indicating that single-trial responses that were more specific to the correct template did not lead to improved target choices.

### Figure 6

From the main results, it seems that single-trial responses are less correlated to the average than a bootstrapped version of it, and that they are only slightly predictive of subsequent behaviour, in only a few brain areas. However, the available information might be more than enough to generate accurate perceptions and behaviour when scaled up to the number of neurons actually involved in the task. To explore this possibility, we first sub-sampled the population of recorded neurons in each brain area at 10 different levels from *N/*10 to *N* . We then extrapolated how metrics like the Specificity Index would evolve as the number of available neurons grew. Stimulus-specificity tended to grow with sample size (Fig. 6A) for Data set 2 but not for Data set 1, raising doubts for the argument that maybe in a realistic population sampled by a downstream neuron (e.g. 30.000 inputs), template matching would be quite strong. Indeed even if the Specificity Index increases in Data set 2, the rate of change is not remarkable. However, as we can see in the bottom part of that plot, Ω did increase for both data sets, and the rate of change is substantial for some areas in Data set 2 (and for both areas in Data set 1). In other words, the single-trial match to the average template may become more indicative of subsequent behavioural choices with larger neuron numbers, even if they are not more stimulus-specific.

### Supplementary Figure 8

Together, the template-matching results for Data set 2 suggest that the relation between single-trial population responses and their trial-averaged response templates is both less strong and less stimulus-specific than what one would expect from a down-sampled representation of the average. Most importantly, single-trial responses that better resembled the correct time-averaged template did not evoke better target choices. One possibility is that ‘average template matching’ happens in a multi-dimensional way. To take a first step at exploring this possibility, we repeated the analyses shown in Fig. 5 by characterizing population responses using Principal Component Analysis (PCA) via Singular Value Decomposition (SVD), and quantifying their resemblance to an average template in this dimensionally-reduced space (Fig. S8).

Generally, the resulting single-trial response vectors did overall not represent their corresponding average vectors more effectively than the Pearson correlation averages we had explored previously. In PCA space, single-trial vectors matched average vectors more closely than would be predicted from bootstrapping (Fig. S7A), but the match was nevertheless weak: the distance between a single-trial vector and its corresponding average template was typically 5 − 10 times larger than the distance between correct and incorrect template. Consistently with this, the specificity of single-trial responses for the correct average template was low, even more so than for linear correlations (Fig. S7C; t-test for difference from Zero: *n* = 90 to 3123; *t* = 0.6 to 17.0; *p <* 0.01 except for *p*(*EPd*) = 0.04 and *p*(*SI*) = 0.56, corrected for multiple comparisons using a FDR procedure with a family-wise error rate of *α* = 0.05). Specificity indices were clustered tightly around zero. Given that PCA vector distances are not upper-bounded to 1 and often took on values between 5 and 10 (Fig. S8A), specificity indices *<* 1 imply negligible differences between the single-trial distances to correct and incorrect templates S8B. Single-trial responses that were more similar and/or specific to the correct average vector also resulted in only slightly more correct behavioural choices (Fig. S8C-E; Mann-Whitney’s U-test for differences in single-trial distances in hit and miss trials: *n* = 90 to 3123, Ω between 0.38 and 0.55; all *p <* 0.01; Mann-Whitney’s U-test for differences in single-trial specificity in hit and miss trials: *n* = 90 to 3123, Ω between 0.5 and 0.61; all *p <* 0.01; corrected for multiple comparisons). Overall, these results demonstrate that just like for Pearson correlations, resemblance of single-trial to average vectors in PCA space did not seem to drive neuronal processing in a decisive way across most brain areas. However, since PCA is a linear method too, this still leaves open the possibility that non-linear methods may reveal accurate template matching of single trials.

## SUPPLEMENTARY FIGURES

**FIG. S1.**
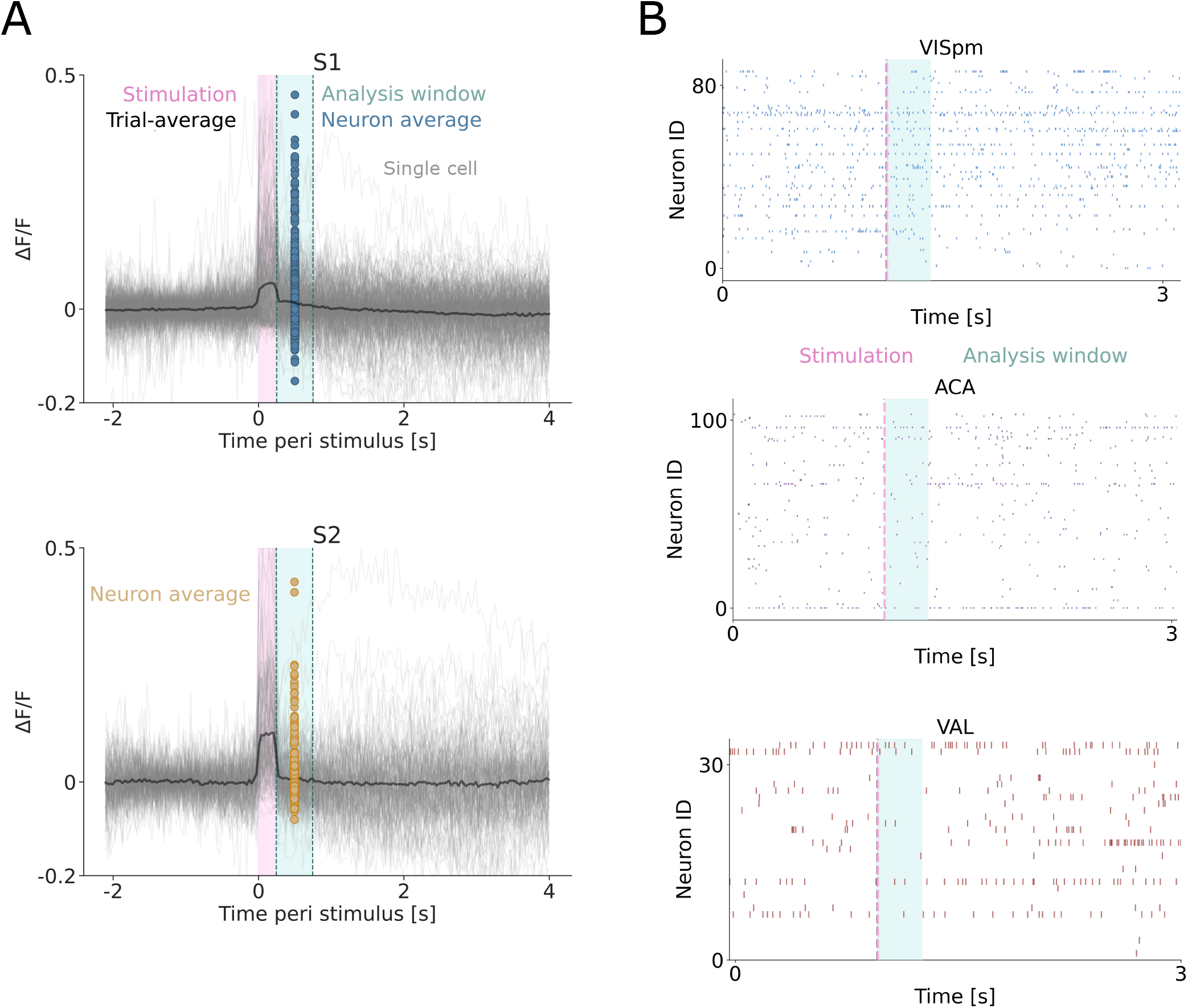
Example of neuronal responses for both Data sets. A) For Data set 1, we show a session, for all neurons over time, aligning the responses to stimulus onset (pink). Traces of individual cells are depicted in light gray, their trial-average in black. In cyan, we show the analysis window (500*ms*), within which we take the time-average (colored dots; blue for S1, orange for S2) that we use to construct the population vectors we used for the analyses. B) Spike traces for different representative areas in Data set 2 (a stimulus informative one [blue, primary visual cortex, VISpm], a choice-informative one [red, ventral anterior-lateral complex of the thalamus, VAL] and a both-informative one [purple, anterior cingulate area, ACA]). Each one comes from a different session and has a different number of neurons. As before, in pink we show the stimulus presentation and in cyan the analysis window (200*ms*) that we used to construct the population vectors in the analyses.

In addition, by pooling stimulus pairs with large and small contrast differences into just two stimulus categories - ‘target left’ and ‘target right’ - we may have caused the resulting average templates to appear less distinctive. Specifically, difficult stimulus pairs might ‘blur the boundaries’ between average templates. To estimate to what extent the poor specificity of single-trial responses in Data Set 2 was caused by difficult stimulus conditions, we computed Specificity Index and Behavioural Relevance separately for difficult and easy trials. Both Specificity and Relevance increased significantly in a majority of brain areas when only taking into account stimulus pairs with large contrast differences (see Fig. S8). This suggests that in Data Set 2, average population firing rates were both specific and behaviourally relevant on a single-trial level when processing coarse stimulus information. In contrast, average population firing rates were neither specific not behaviourally relevant when animals were engaged in finer contrast discrimination. In this context, it is important to note that animals were also highly successful in discriminating these difficult stimulus pairs. This suggests that fine contrast discrimination relied on other coding modalities than average firing rates.

**FIG. S2.**
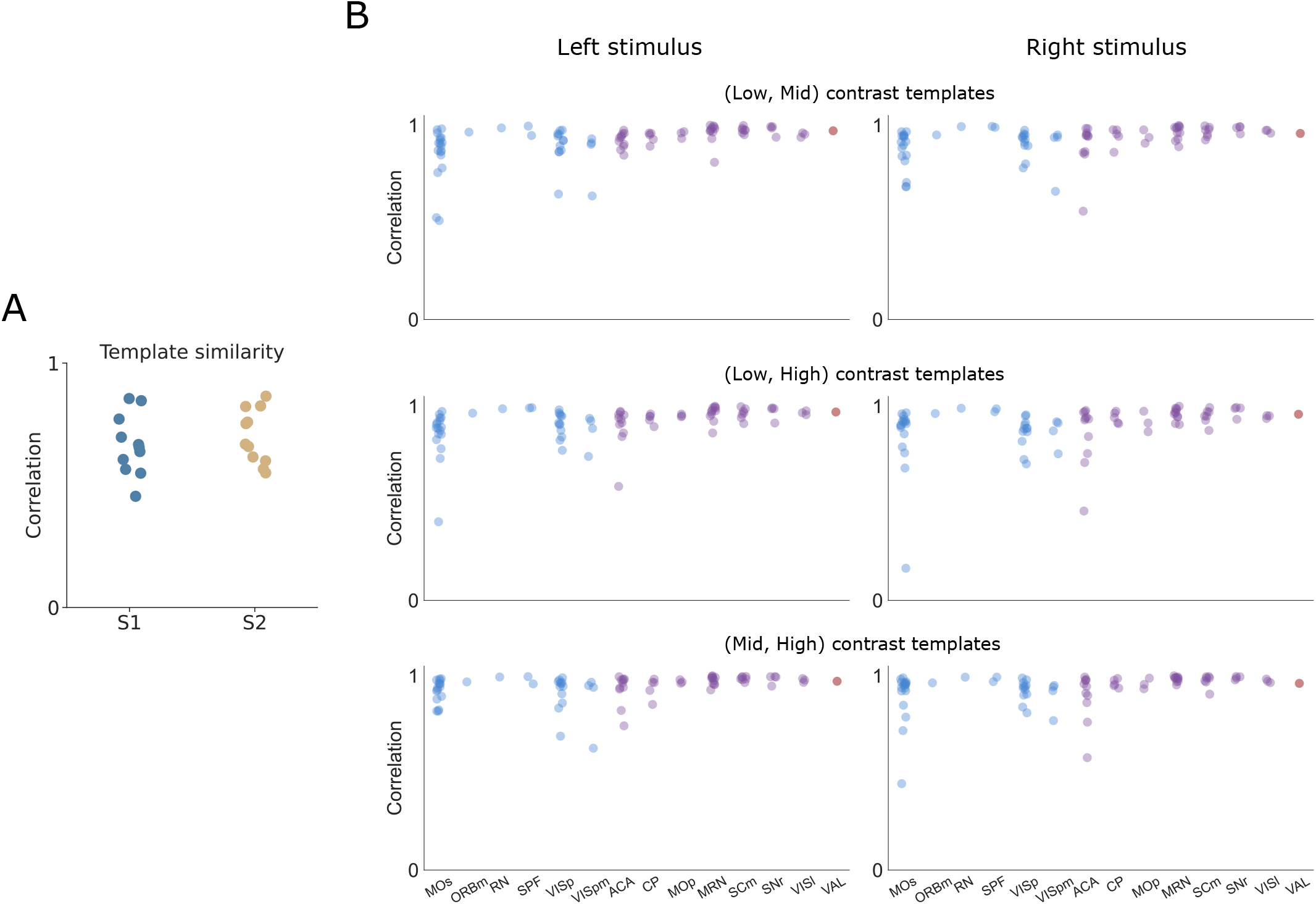
Correlations between response templates for different stimulus constellations. A) For Data set 1, we pool together those trials with a low number of stimulated neurons (*n* ∈ [5, 20]) and compare the trial-averaged response with those trials with a higher number of stimulated neurons (*n* ∈ [30, 50]). Their correlation generally exceed 0.6, suggesting that one template should be sufficient to represent different contrast constellations. B) For Data set 2, we compute the similarity between templates (for each contrast level: 0.25, 0.5, 1, referred to as low, mid and high, respectively) for each screen (left, right). Their correlation is generally above 0.8, so grouping them together should provide a coherent representation.

**FIG. S3.**
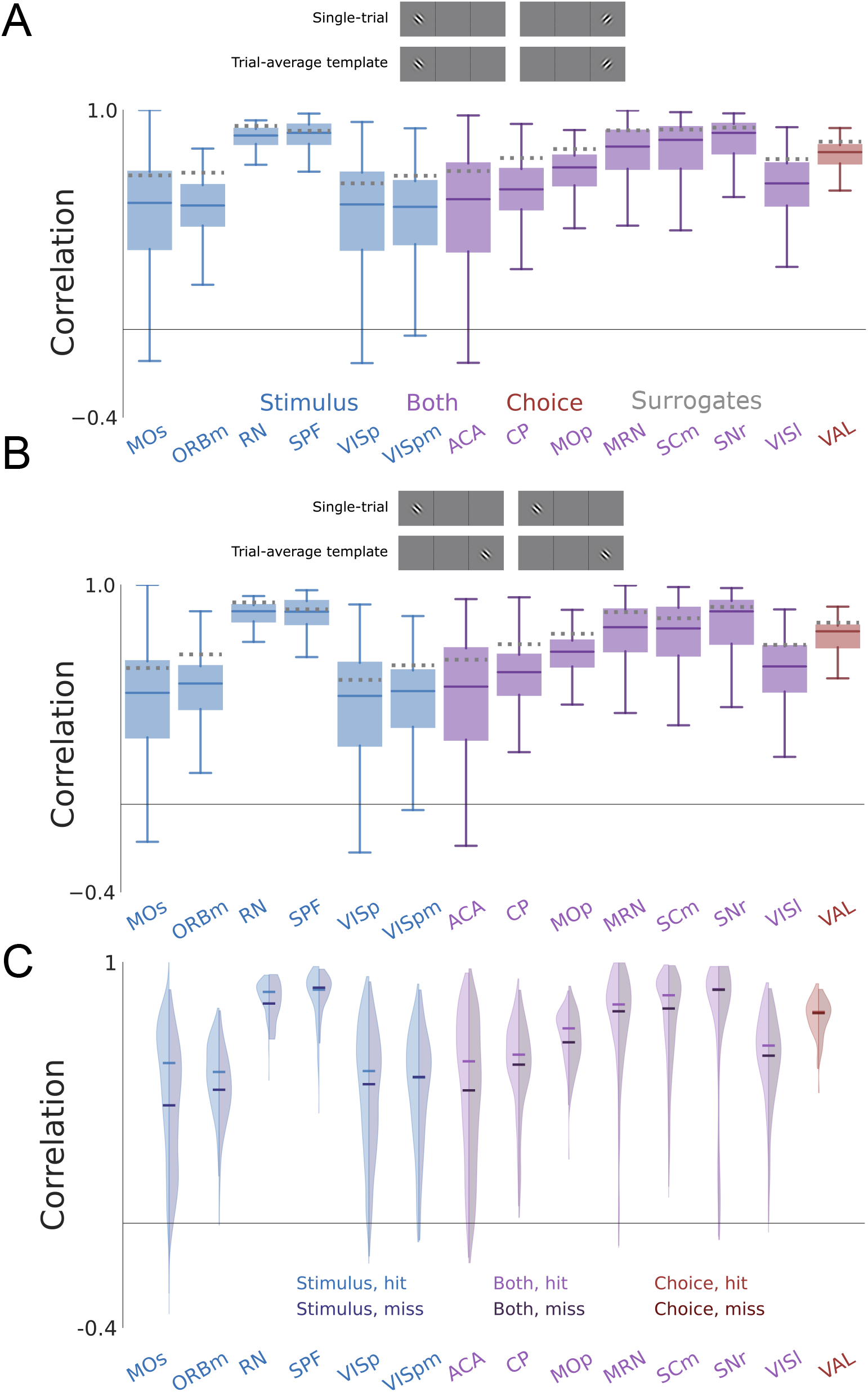
Single-trial responses are not stimulus-specific for Data set 2. A) Distribution of the correlations between single-trial responses and the matching trial-averaged response templates. The top diagram shows the criterion to “match” single-trial responses with the trial-average. Box: 25^*th*^ and 75^*th*^ percentile. Center line: median. Whiskers: 10^*th*^ and 90^*th*^ percentile. Dotted lines: median of bootstrapped data. B) Same as A) but for the non-matching combinations. C) Distribution of the correlations between single-trial responses and the matching trial-averaged response templates, split by behavioural outcome. It can be seen that these distributions are practically identical for all regions.

**FIG. S4.**
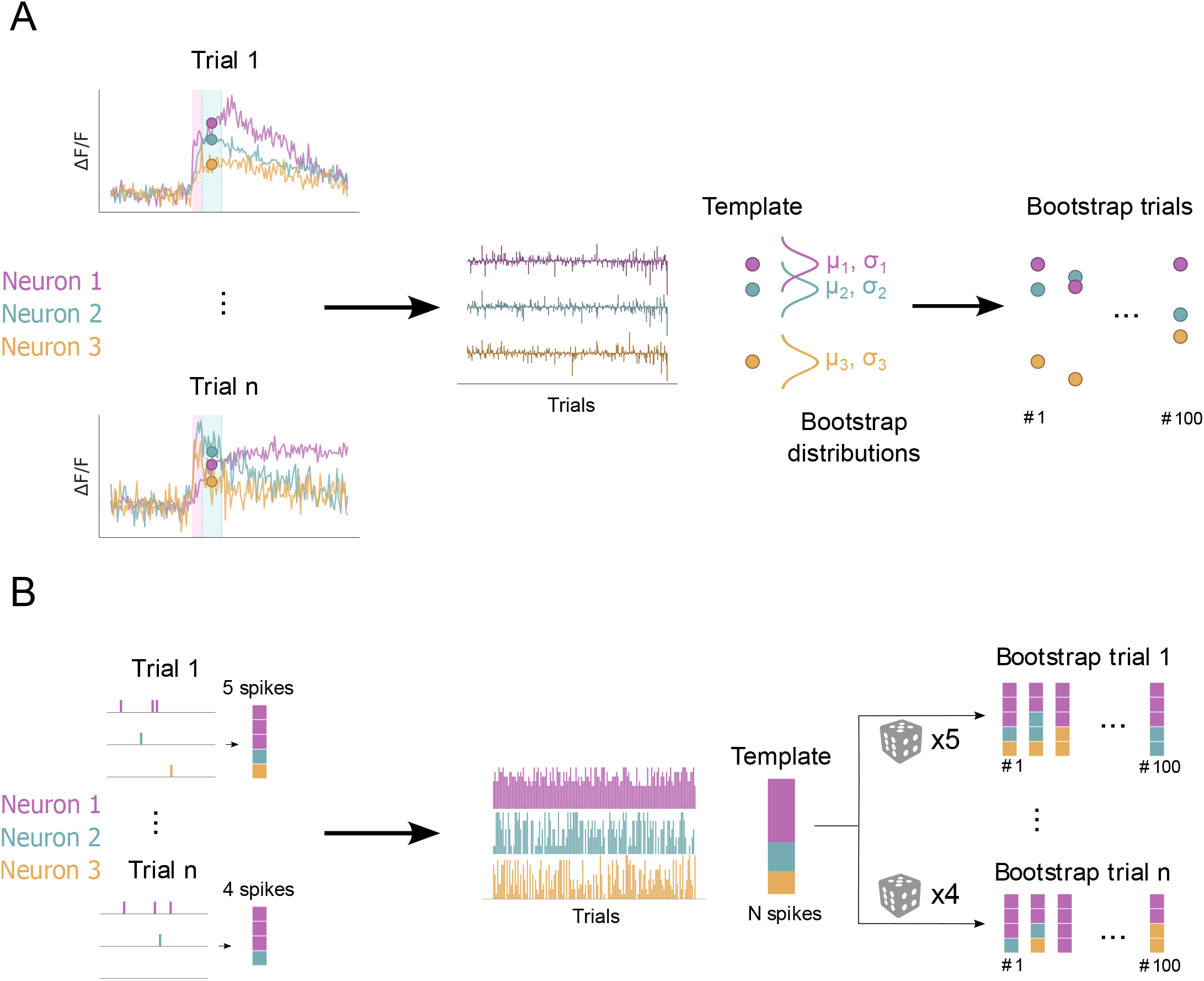
Representation of the bootstrapping procedures. A) In the first data set, we created the templates based on the time-average for each neuron, for each trial. We then constructed a Gaussian distribution centered around the trial-average and with a spread equal to the trial-variance. We repeated this for each neuron and then sample 200 times for each stimulus set. B) For data set 2, we fixed the number of spikes on each trial, but randomly assigned to a neuron with a probability according to how often it spiked in the average template. We repeated this procedure 100 times.

**FIG. S5.**
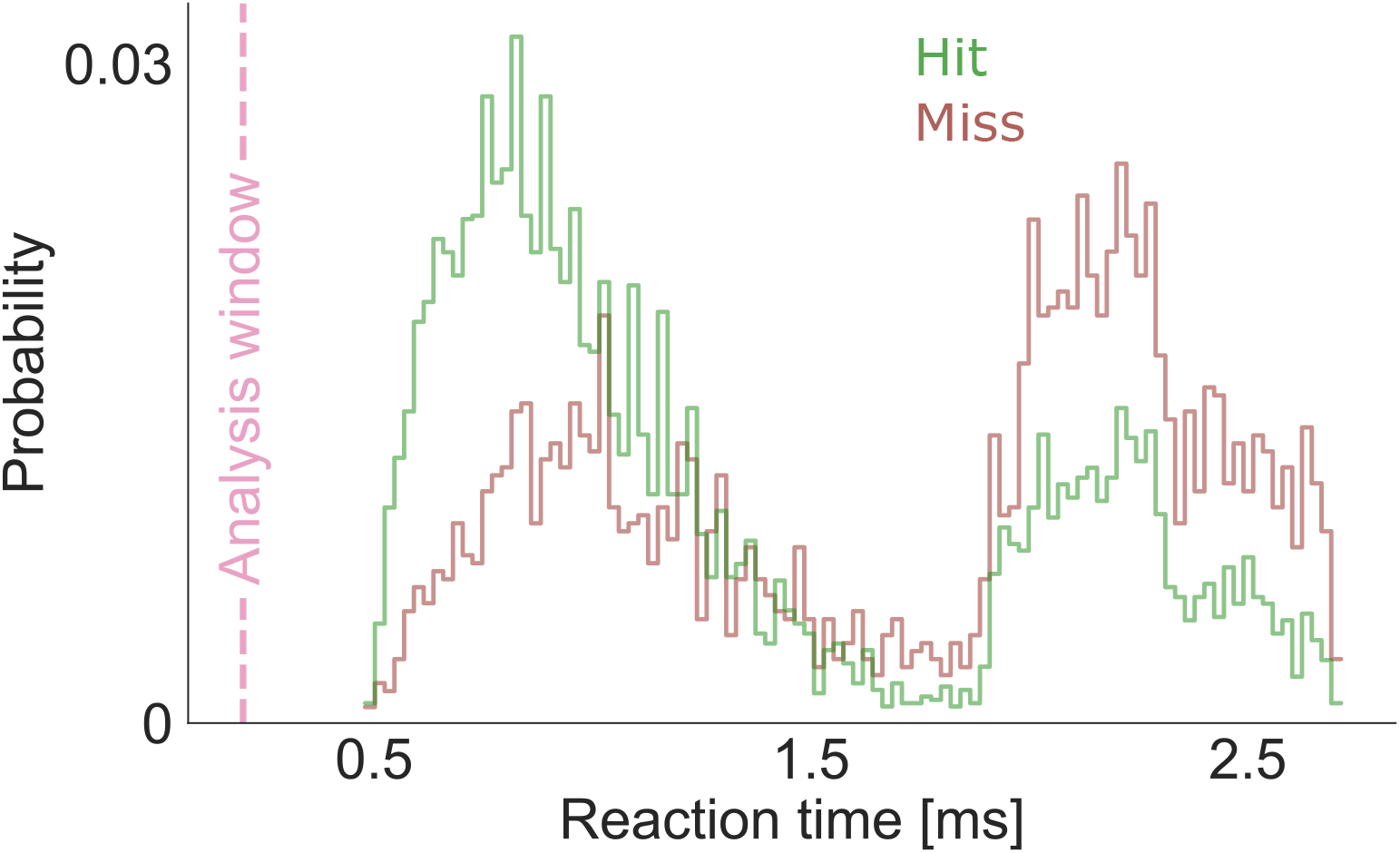
Reaction time distribution. General distribution for reaction times, over all sessions, split by hits and misses. Consistent with the literature, misses significantly (Mann-Whitney’s U-test, *p* = 1.88 * 10^*−*172^, *A* = 0.68) imply longer reaction times. The vertical line indicates the width of the time window in which we have performed all of our analyses (200*ms*). We have selected our analysis window of because of its likely relevance to stimulus processing and behavioural decision making.

**FIG. S6.**
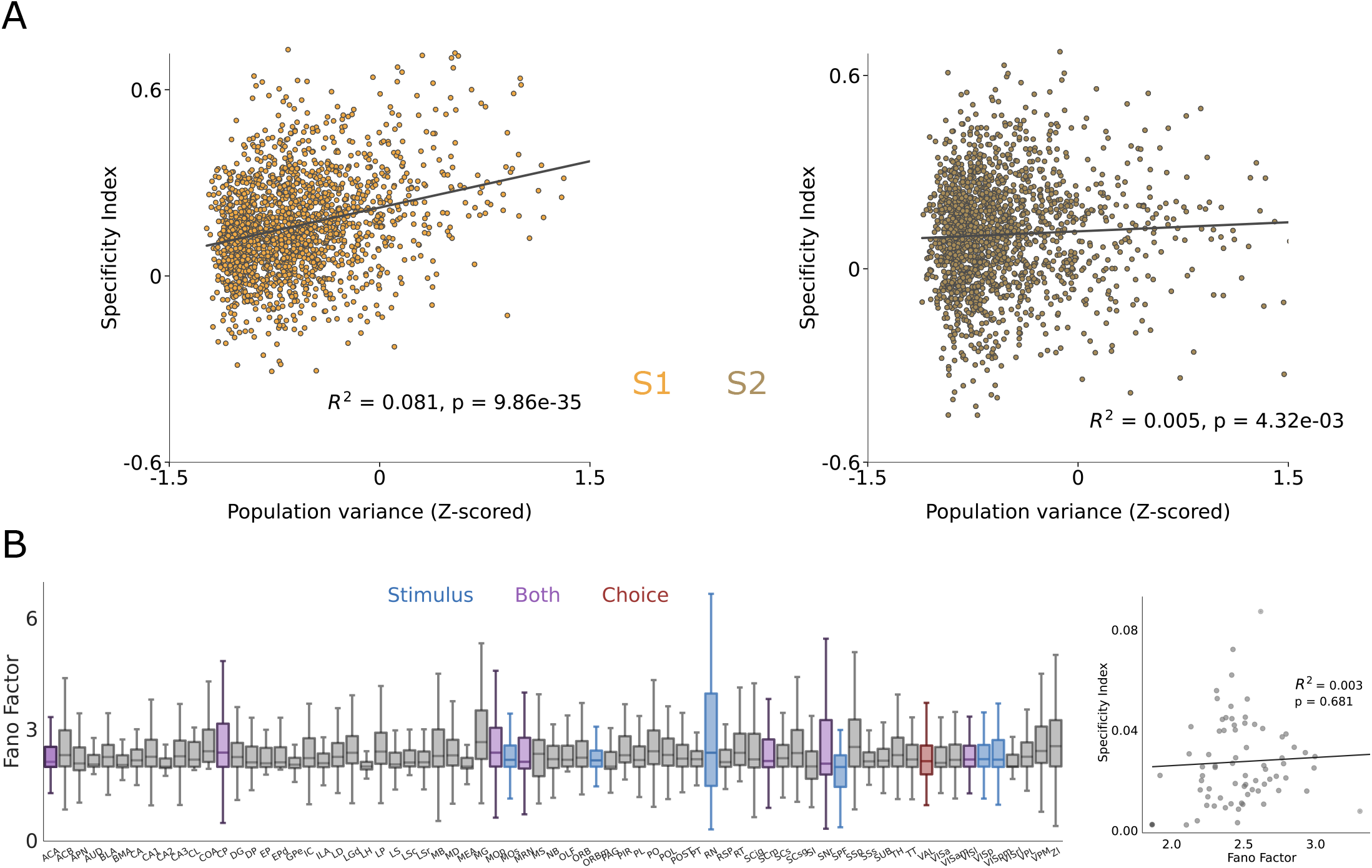
Population variability is not predictive of stimulus-specificity. A) Scatter of the Specificity Index for different population variances. There is virtually no correlation for both regions. B) Fano factor distribution over all recorded regions. As for Data set 1, variability is uncorrelated with the Specificty Index and it is, thus, not capturing the same effect.

**FIG. S7.**
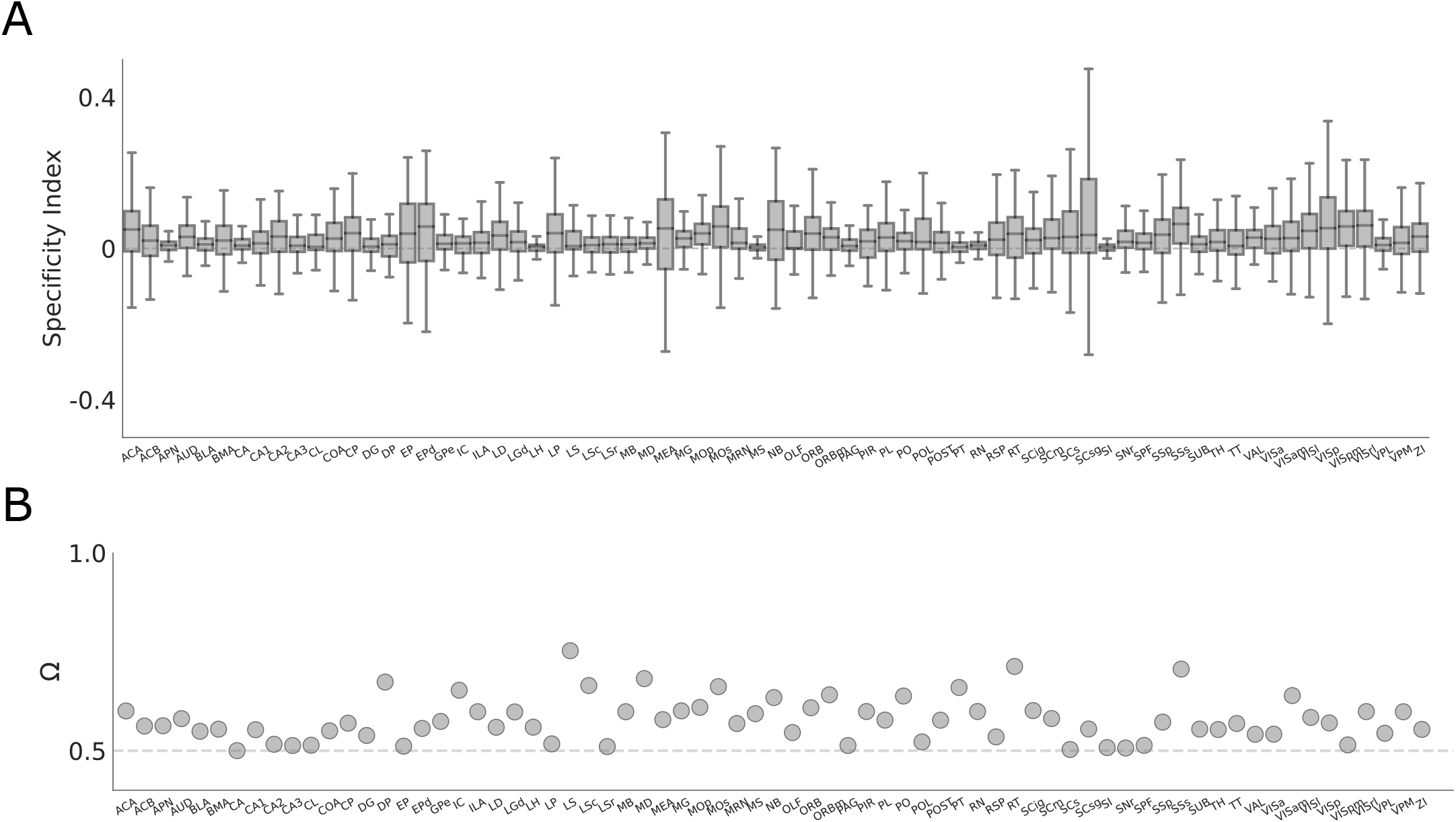
Specificity Index and Behavioral Relevance for all areas. Same as Fig. 5B,E, but without pre-selecting informative areas. Results are preserved in general: across areas, single-trial responses are barely stimulus-specific (Specificity Indices are tightly clustered around 0, with a median value of 0.019) and not very behaviourally relevant (median value of Ω = 0.57).

**FIG. S8.**
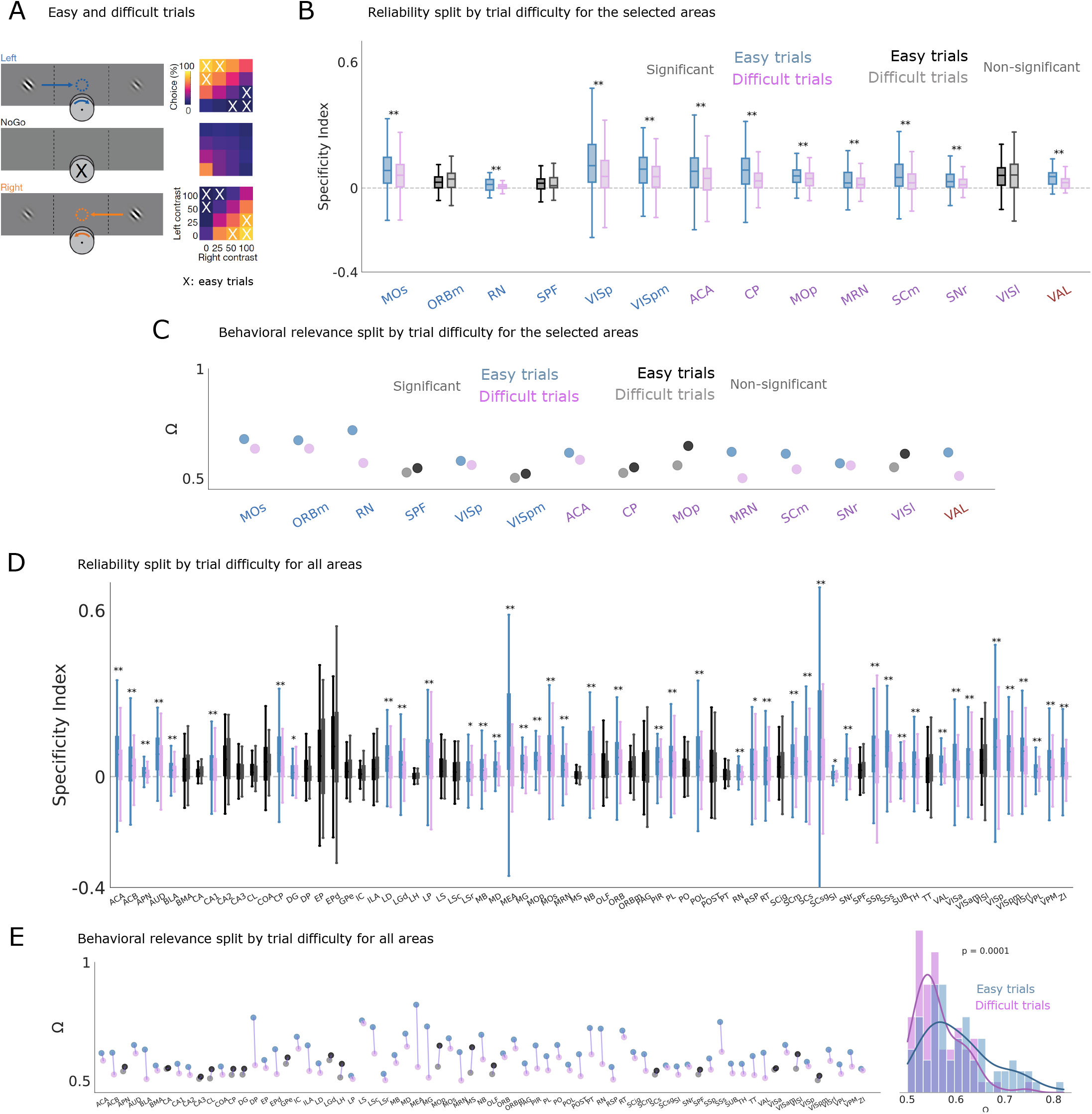
Easy trials are more stimulus-specific and behaviourally relevant than difficult trials for Data set 2. A) Task structure and behavioural proficiency; easy trials are taken to be the extremal cases: low contrast shown on one screen and a high one in the other. B) Distribution of the Specificity Index for easy and difficult trials, over the selected areas. In color, significant comparisons. Box: 25^*th*^ and 75^*th*^ percentile. Center line: median. Whiskers: 10^*th*^ and 90^*th*^ percentile. Dotted lines: median of bootstrapped data. C) Behavioural relevance (Ω) for easy and difficult trials, over all selected areas. D) Same as B) but for all recorded areas. E) Same as C), but for all recorded areas. Average responses are more behaviourally relevant for easy than for difficult trials (right).

**FIG. S9.**
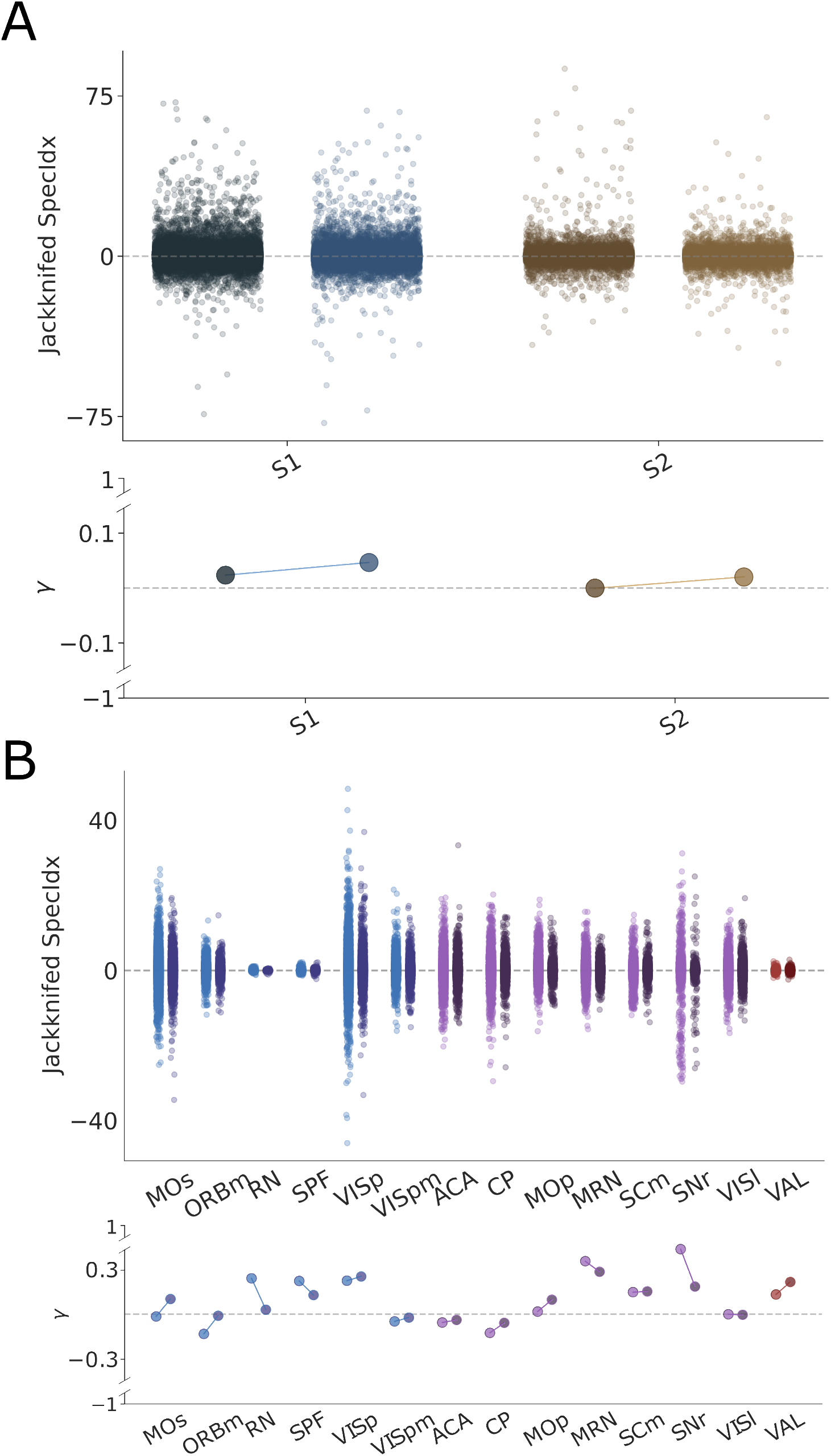
Jackknife analyses. We remove one neuron at a time and compute the impact in the Specificity Index, calculated as 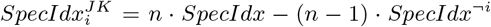, for neuron *i*. We show the symmetry (*γ*) of each distribution around 0 using Yule’s coefficient (Methods). This measure is bounded between −1 and 1, with each of them meaning completely skewed towards negative or positive values, respectively. A) Change in single-trial correlations to the correct average template when one individual neuron was removed. Data points: Trials. Colors: see inset legend. B) Same for single-trial specificity.

**FIG. S10.**
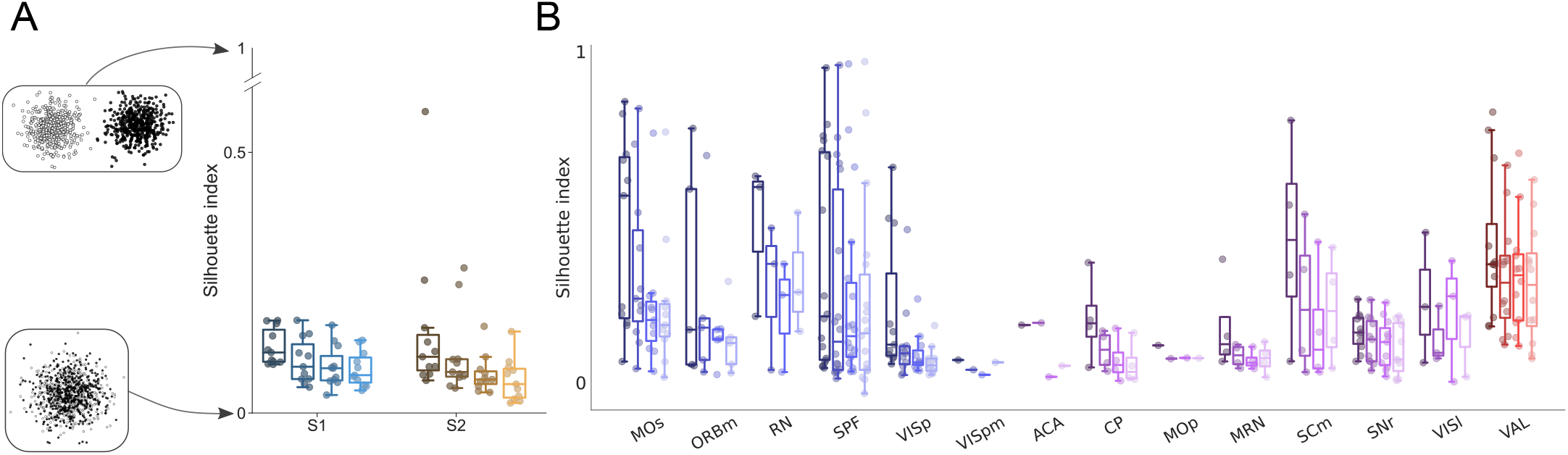
Unsupervised trial-clustering for both data sets. Silhouette Index (SI) for both Data sets. This quantity measures cluster compactness (see Methods), with 0 indicating complete overlap between spread clusters and 1 meaning perfectly separated ones. Different colors represent a different number of clusters, from *k* = 2 to *k* = 5. A) SI for Data set 1. In this case, clusters are not compact and they are better distinguishable (for both brain areas) when *k* = 2. B) SI for Data set 2. Clusters are more compact than in the previous case, with *k* = 2 also being the best option. Thus, for analyses in the main text, we decided to group trials into two clusters.

**FIG. S11.**
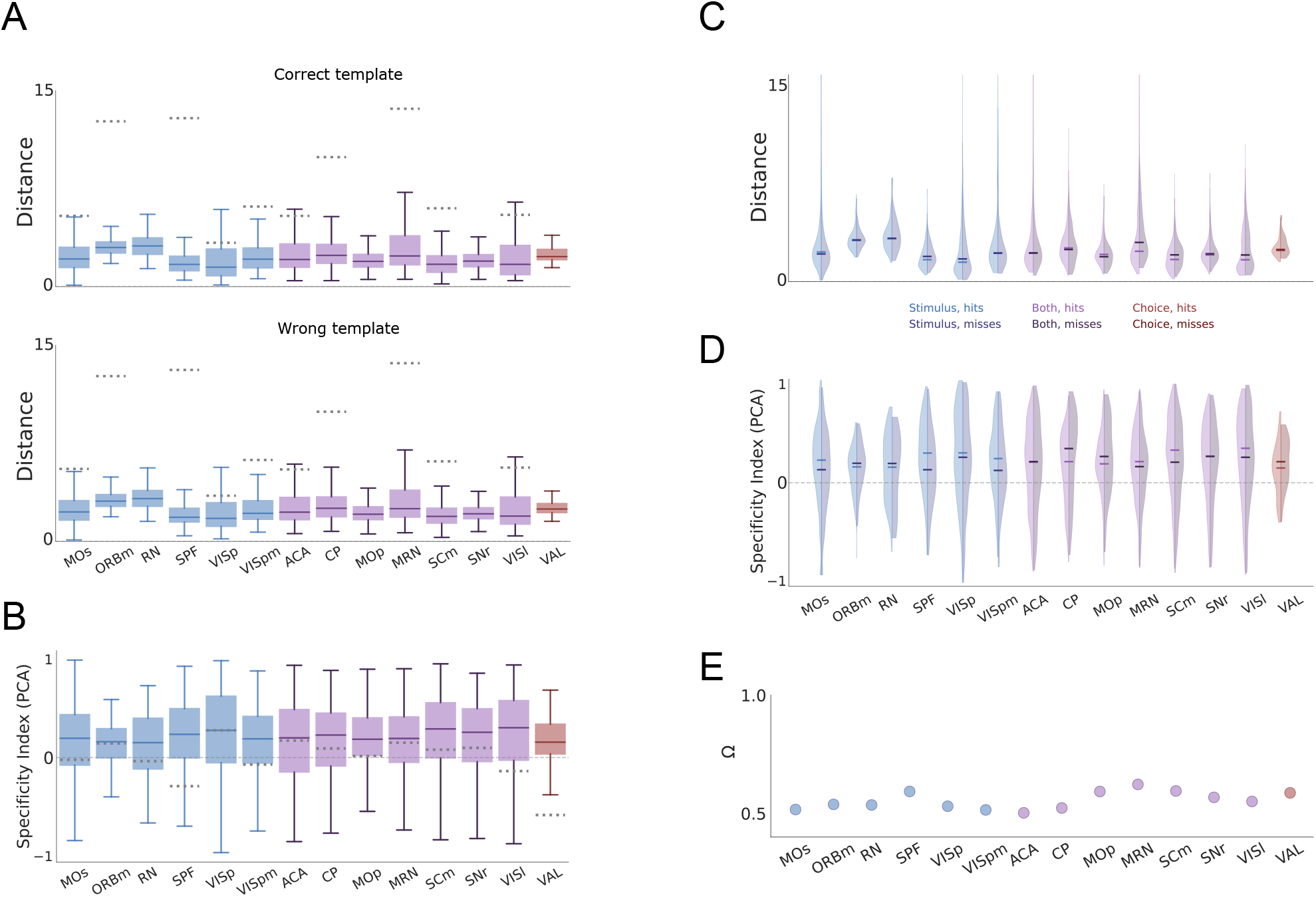
PCA is not helpful in making single-trial responses more stimulus-specific or behaviorally relevant for Data set 2. A) Distribution of single-trial normalized distances in PCA space (Methods) between response vectors and the trial-averaged response template for the correct (top) and wrong (bottom) stimulus constellation. B) Same as B, but the Specificity index of single-trial responses across brain areas, defined as the difference between the normalized distance to the wrong and correct template. Solid gray line highlights the Specificity Index of 0.0, which translates to exactly equal correlation to correct and wrong template. Dotted lines represent the specificity index of the medians of the bootstrapped values for each recorded area. C) Same as A (top), split by hits and misses. These distributions almost completely overlap, for all brain areas. D) Same as B, split by hits and misses. As in the previous panel, these distributions are practically coincident. E) Behavioral Relevance Index for all brain areas. They range from 0.50 to 0.62, with Ω = 0.5 meaning perfect overlap.

**FIG. S12.**
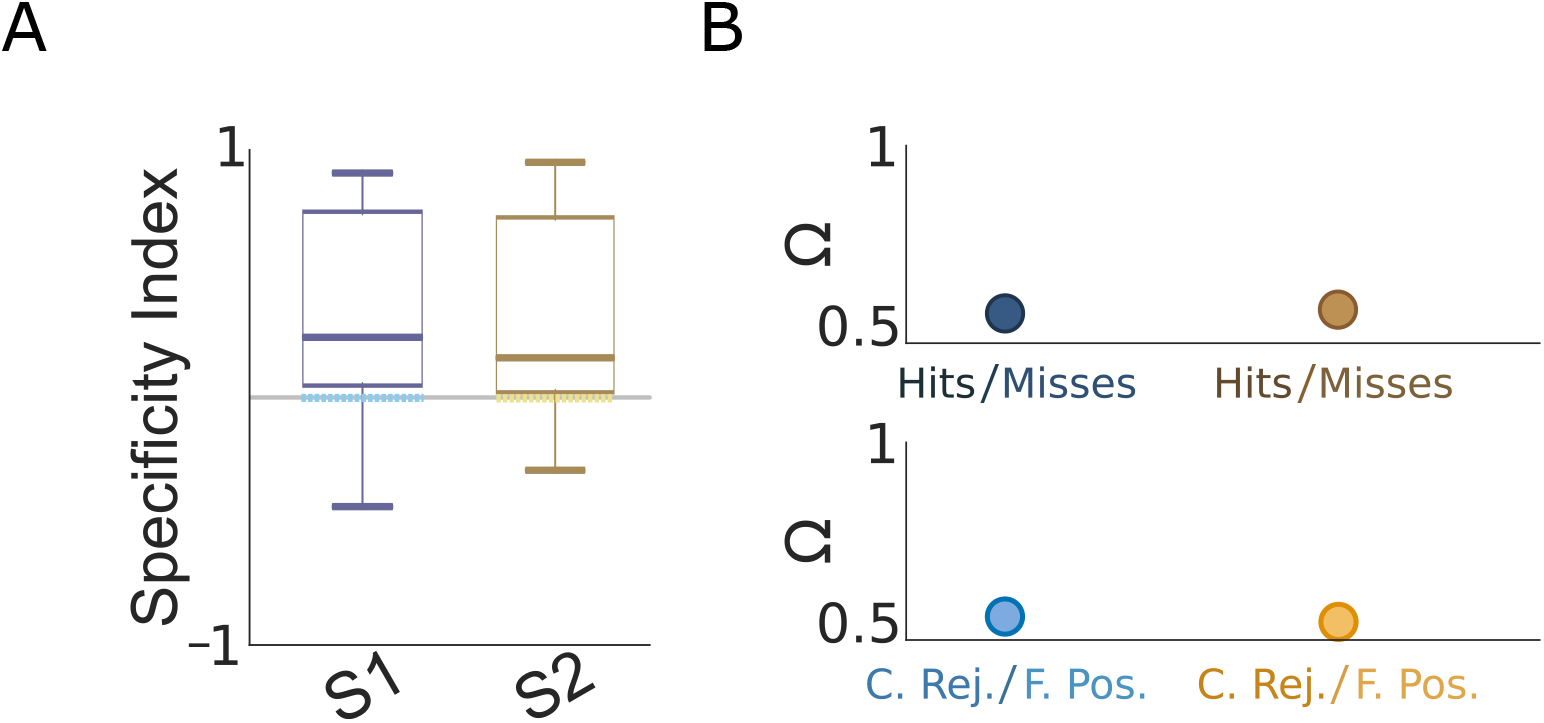
PCA is helpful in making single-trial responses more stimulus-specific but heavily reduces behaviorally relevant for Data set 1. A) Same as Fig. S8B, for Data set 1. B) Same as Fig. S8E, for Data set 1. In this case, the Specificity Index increased modestly (from 0.13 to 0.21) at the expense of severely hindering behavioral relevance (from 0.83 to 0.52).

**FIG. S13.**
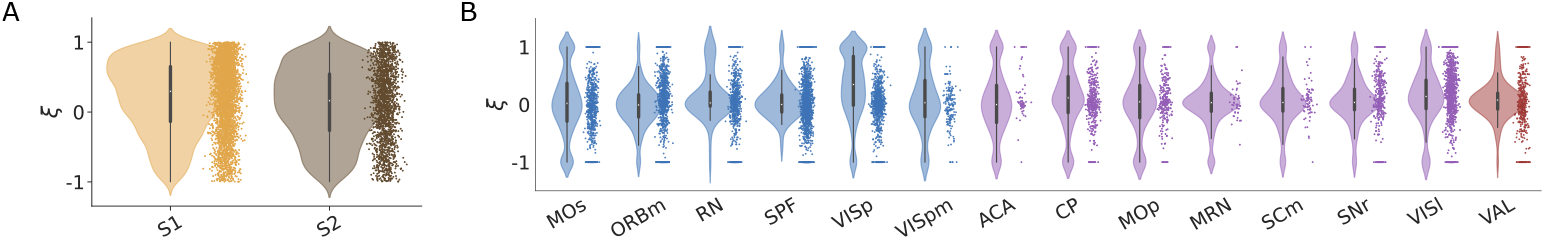
Selectivity distributions for both Data sets..

